# A SUMO-dependent feedback loop senses and controls the biogenesis of nuclear pore subunits

**DOI:** 10.1101/188441

**Authors:** Jérôme O. Rouvière, Manuel Bulfoni, Alex Tuck, Bertrand Cosson, Frédéric Devaux, Benoit Palancade

## Abstract

While the activity of multiprotein complexes is crucial for cellular metabolism, little is known about the mechanisms that collectively control the expression of their components in response to the cellular demand. Here, we have investigated the regulations targeting the biogenesis of the nuclear pore complex (NPC), the macromolecular assembly mediating nucleocytoplasmic exchanges. Systematic analysis of RNA-binding proteins interactomes, together with *in vivo* and *in vitro* assays, revealed that a subset of *NPC* mRNAs are specifically bound by Hek2, a yeast hnRNP K-like protein. Hek2-dependent *NPC* mRNAs translational repression and protein turnover are further shown to finely tune the levels of NPC subunits. Strikingly, Hek2 binding to its target mRNAs requires prior desumoylation by the NPC-associated SUMO protease Ulp1. Mutations or physiological perturbations altering NPC integrity lead to a decrease in the levels of active Ulp1 and to the accumulation of sumoylated, inactive forms of Hek2. Our results support the existence of a quality control mechanism involving Ulp1 as a sensor of NPC integrity and Hek2 as a repressor of NPC biogenesis.

## INTRODUCTION

Virtually all cellular processes rely on the function of multiprotein assemblies. While their stoichiometry has to be tightly controlled to prevent an imbalance of subunits that could interfere with their assembly or titrate their targets, their global abundance has also to be adjusted in response to the cellular demand (Harper and Bennett, 2016). Multiple layers of mechanisms have been reported to partake in the accurate biogenesis of multisubunit complexes. First, all the steps in the gene expression pathway, including mRNA synthesis, processing, transport, stability and translation, can be regulated in a coordinate manner, either to lead to the proportional synthesis of the different subunits of multiprotein assemblies, a prominent strategy in prokaryotes (Li et al., 2014), or to respond to environmental or physiological cues, as exemplified by the ribosome biosynthesis pathway (Lempiainen and Shore, 2009). In this frame, a pivotal role has emerged for transcriptional regulators and RNA-binding proteins, the latter being in particular capable to tune the translation rate of their target messenger ribonucleoparticles (mRNPs). Second, molecular chaperones and assembly factors can further assist the assembly of multiprotein complexes, as also described for ribosomes (Lempiainen and Shore, 2009), in some cases in a cotranslational manner (Natan et al., 2017). Finally, excess complexes or unassembled, orphan polypeptides can be targeted for degradation by the proteasome or the lysosome (Wolff et al., 2014), these quality control processes being critical to adjust stoichiometry and to cope with altered protein dosage (Dephoure et al., 2014; McShane et al., 2016). However, despite our improved knowledge in proteome dynamics, the specific mechanisms at play for most multiprotein complexes remain largely unknown.

The nuclear pore complex (NPC) provides a paradigmatic example of an essential multisubunit complex whose homeostasis is crucial yet poorly understood. NPCs are megadalton-sized proteinaceous assemblies embedded at the fusion points of the nuclear envelope and formed of modular repeats of ∼30 distinct protein subunits - the nucleoporins (Nups) -, which assemble within subcomplexes and organize with a 8-fold rotational symmetry (Beck and Hurt, 2017). The major task of NPCs is the selective nucleocytoplasmic transport of macromolecules, i.e. proteins and RNA-containing particles, a process involving dynamic interactions between the cargoes-transport factors complexes and the phenylalanine-glycine (FG) repeats-harboring nucleoporins that lie within the central channel and the peripheral extensions of the NPC (Floch et al., 2014). The stepwise assembly of nucleoporins to build complete NPCs proceeds through defined pathways, either following mitosis in conjunction with nuclear envelope reformation, or during interphase, the unique assembly mode compatible with the closed mitosis of fungi. Nucleoporins themselves are essential players in NPC assembly, either through scaffolding, or by mediating interactions with chromatin and/or membranes. In addition, non-NPC factors, such as membrane bending proteins, also contribute to NPC biogenesis (Weberruss and Antonin, 2016). While multiple studies have depicted the choreography of NPC assembly, together with their structural organization, little is known about the mechanisms that sustain the timely production of stoichiometric amounts of Nups and that could possibly sense and adjust NPC biogenesis depending on cell physiology.

The high connectivity observed between NPCs and several biological processes could place them in a strategical position to communicate their status to the cell. Indeed, NPCs have been described to contribute to multiple aspects of transcriptional regulation, genome stability and cell cycle progression (Floch et al., 2014). In some situations, these connections are mediated by physical interactions between NPCs and enzymes of the small ubiquitin-related modifier (SUMO) pathway (Palancade and Doye, 2008). Sumoylation is a post-translational modification that can modulate the binding properties or the conformation of its targets, ultimately impacting their stability, their localization or their biological activity (Flotho and Melchior, 2013). Among the distinct enzymes of the sumoylation/desumoylation machinery shown to associate with NPCs, the conserved SUMO protease Ulp1 has essential functions in SUMO processing and deconjugation in budding yeast. The docking of this enzyme to the nucleoplasmic side of NPCs is essential for viability (Li and Hochstrasser, 2003; Panse et al., 2003) and is believed to involve its nuclear import through karyopherins, followed by its association with several nucleoporins (Zhao et al., 2004; Lewis et al., 2007; Palancade et al., 2007; Srikumar et al., 2013; Hirano et al., 2017). Proper NPC localization of Ulp1 has been shown to be critical for the spatio-temporal control of the sumoylation of certain targets, some of them being important for genetic integrity or gene regulation (Li and Hochstrasser, 2003; Makhnevych et al., 2007; Palancade et al., 2007; Texari et al., 2013).

Here, we report an original mechanism by which the synthesis of NPCs subunits is regulated in response to changes in NPC integrity in budding yeast. We show that a subset of Nups-encoding mRNAs is defined by the specific binding of the translational regulator Hek2. Hek2-regulated *NPC* mRNAs translation and protein turnover are further shown to finely tune the levels of the corresponding nucleoporins. Strikingly, Hek2 binding to *NPC* mRNAs is prevented by sumoylation, a process reversed by the SUMO protease Ulp1. Mutant or physiological situations in which NPCs functionality is compromised are associated with the loss of Ulp1 activity and the subsequent accumulation of sumoylated Hek2 versions that are inactive for *NPC* mRNAs translational repression. We propose that Ulp1 and Hek2 are respectively the sensor and the effector of a feedback loop maintaining nucleoporins homeostasis.

## RESULTS

### A unique mRNP composition for a subset of nucleoporin-encoding mRNAs

In order to unravel novel mechanisms regulating NPC biogenesis, we systematically analyzed the association of Nups-encoding (*NPC*) mRNAs with different RNA-binding proteins (RBP) in budding yeast. For this purpose, we took advantage of previously published large-scale datasets obtained through RNA immunoprecipitation (RIP; Hieronymus and Silver, 2003; Kim Guisbert et al., 2005; Hogan et al., 2008; Hasegawa et al., 2008), cross-linking immunoprecipitation (CLIP; Wolf et al., 2010) or crosslinking and analysis of cDNA (CRAC; Tuck and Tollervey, 2013). We collected the association data for 39 *NPC* mRNAs (encoding Nups and NPC-associated proteins, **Fig. 1A** and **Fig. S1A**) with a panel of 10 mRNA-associated factors involved in different stages of mRNA metabolism, including assembly into mRNP (Sto1), processing (Npl3, Nab4/Hrp1), nuclear export (Yra1, Nab2, Mex67), degradation (Xrn1, Ski2, Mtr4) or mRNA localization/translation (Hek2) (**Fig. 1A**). This analysis revealed that *NPC* mRNAs have generally the same typical features of expressed, protein-coding RNAs, e.g. they readily associate with mRNA export factors (Mex67, Nab2), but not with the non-coding RNA degradation machinery (Mtr4) (**Fig. 1A**, bottom right panel). Strikingly however, a small subset of *NPC* mRNAs (namely *NUP170, NUP59, NUP188, NUP116, NUP100, NSP1* and *NUP1*) appeared to specifically bind the conserved <u>He</u>terogeneous nuclear ribonucleoprotein <u>K</u>-like factor Hek2 (a.k.a. Khd1; Irie et al., 2002; Paquin et al., 2007), a feature detected in four independent datasets (**Fig. 1A**, bottom left panel). The enrichment of certain *NPC* mRNAs among Hek2-bound targets appeared significant in a Gene Set Enrichment Analysis (p=0.02) and was neither a mere consequence of the different expression levels of these particular transcripts (**Fig. S1B**), nor a general feature of any multiprotein complexes, since it was not observed when similar analyses were performed for mRNAs encoding proteasome or exosome subunits (**Fig. S1C**).

**Figure 1.**
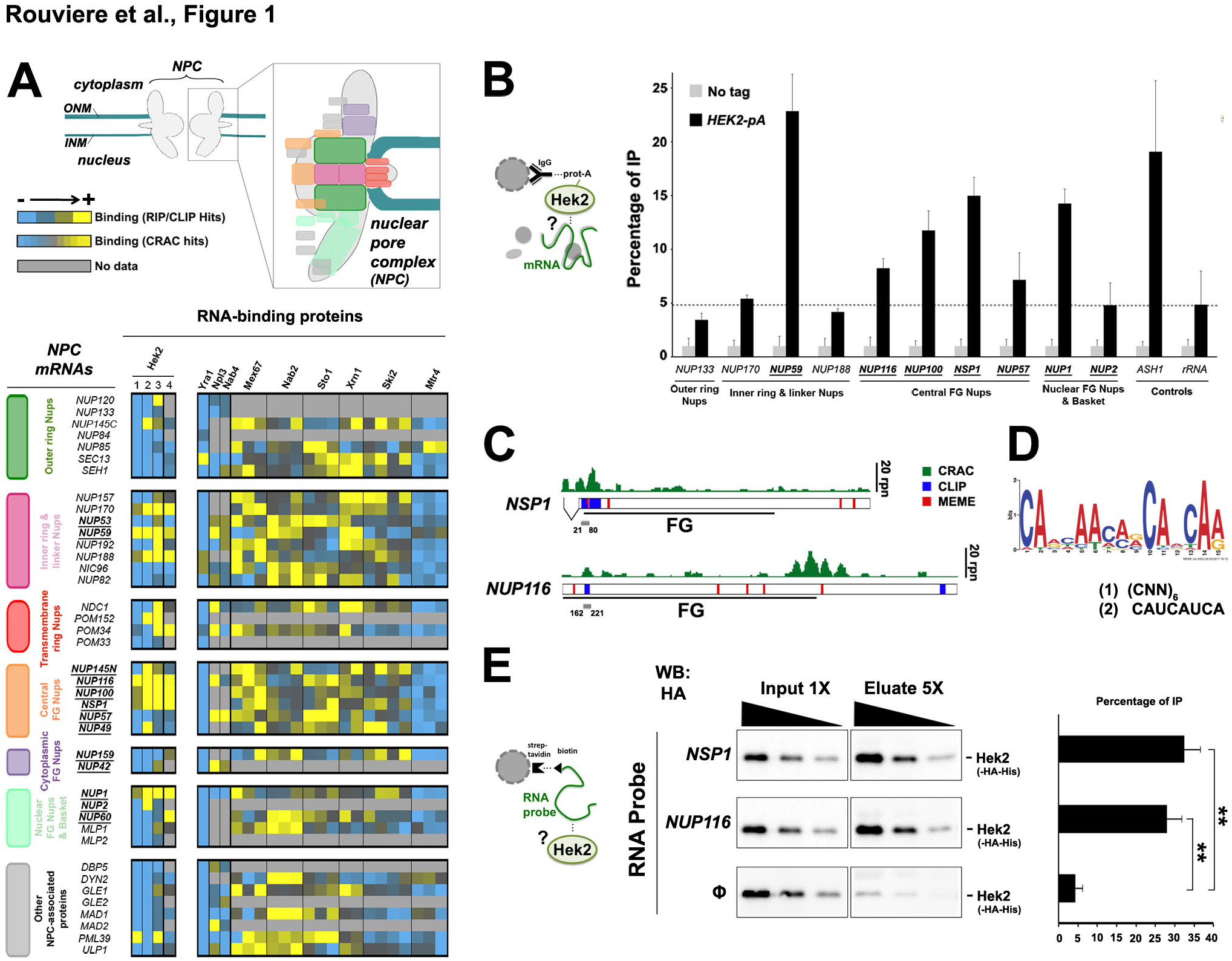
– The hnRNP K-like protein Hek2 specifically associates with a subset of NPC mRNAs. **A,** *Top*, schematic representation of the yeast nuclear pore complex (NPC). ONM, outer nuclear membrane. INM, inner nuclear membrane. NPC subcomplexes are represented as colored boxes. *Bottom*, mRNAs encoding nucleoporins and NPC-associated proteins are sorted by subcomplexes and the strength of their association to the different indicated RNA-binding proteins (RBP) is represented by a color code, as scored in distinct RIP, CLIP or CRAC datasets. Bright yellow indicates the preferred association of a given mRNA to the RBP of interest. For Sto1, Mtr4, Nab2, Mex67, Xrn1 and Ski2, multiple repetitions are displayed (Tuck and Tollervey, 2013). For Hek2, the results from independent studies are represented, as follows: (1) (Hasegawa et al., 2008), (2) (Wolf et al., 2010), (3) (Hogan et al., 2008), (4) (Tuck and Tollervey, 2013). FG-Nups appear in bold, underlined. The *NUP145* mRNA, which gives rise to both Nup145-N and Nup145-C nucleoporins, is displayed for each of the subcomplexes to which these proteins belong. See Methods for details. **B**, Hek2-pA-associated mRNAs were immunopurified and quantified by RT-qPCR using specific primer pairs. Percentages of IP (mean ± SD; n=3) are the ratios between purified and input RNAs, further normalized to the amount of purified bait and set to 1 for the “no tag”. A schematic representation of the assay is shown. **C,** Overview of Hek2 binding sites on *NSP1* and *NUP116* mRNAs. The number of CRAC hits (rpn, reads per nucleotide; Tuck and Tollervey, 2013), the position of CLIP fragments (Wolf et al., 2010) and the occurrences of the binding site found by the MEME analysis are indicated. The positions of the FG-coding region and of minimal Hek2-binding sites used for *in vitro* pull down (in grey) are represented. The broken line indicates the *NSP1* intron. **D,** MEME result from *NUP59*, *NUP116*, *NUP1*, *NSP1* and *NUP100* sequences. (1) and (2) indicate previously identified Hek2-binding sequences (Hasegawa et al., 2008; Wolf et al., 2010). **E**, *Left*, Schematic representation of the assay. Recombinant HA-tagged Hek2 was incubated with streptavidin beads either naïve (Φ) or previously coated with biotinylated RNA probes encompassing Hek2-binding sites from *NSP1 (21-80)* or *NUP116 (162-221). Center*, Decreasing amounts of input and eluate fractions were loaded to allow quantification. *Right*, Percentages of IP (mean ± SD; n=3) are the ratios between Hek2 amounts in the eluate and in the input fractions. ** P &#x003C; 0,01 (Welch’s t-test). **See also Fig. S1**.

To further validate this finding *in vivo*, we immunoprecipitated a protein A-tagged version of Hek2 from yeast cells and analyzed its interaction with *NPC* mRNAs by RT-qPCR. In agreement with our previous findings, Hek2 preferentially associated with *NUP59, NUP116*, *NUP100*, *NSP1* and *NUP1* mRNAs (**Fig. 1B**), to a similar extent as its prototypal target *ASH1* (Irie et al., 2002; Paquin et al., 2007), but not with *NUP133*, *NUP57*, or *NUP2* mRNAs (**Fig. 1B**), for which Hek2 binding was in the same range as its reported, unclear association to rRNA (Tuck and Tollervey, 2013). Preferential binding to *NUP170* and *NUP188* mRNAs was not confirmed, the previous finding from genome-wide studies possibly reflecting their different expression levels in other genetic backgrounds. In contrast, immunoprecipitation of Hpr1, a subunit of the mRNP packaging THO complex, did not reveal any similar preferred association to a subset of *NPC* mRNAs (**Fig. S1D**).

We then asked whether Hek2 was directly associating to this subset of *NPC* mRNAs (i.e. *NUP59*, *NUP116*, *NUP100*, *NSP1* and *NUP1)*, as expected from CLIP/CRAC studies (Wolf et al., 2010; Tuck and Tollervey, 2013). To this aim, we first delineated Hek2-binding sites on these mRNAs by mining CLIP/CRAC data (**Fig. 1C**) and by searching their sequences for common motifs using the MEME software (**Fig. 1C-D**). This *in silico* approach revealed that these mRNAs share a common CA-rich motif (**Fig. 1D**), similar to the two previously reported Hek2 binding sites, i.e. (CNN)_6_ (Hasegawa et al., 2008) and CAUCAUCA (Wolf et al., 2010). As anticipated from a previous study (Wolf et al., 2010), this motif was overlapping some but not all *in vivo* Hek2 binding peaks as defined by CLIP or CRAC, allowing us to define putative minimal bound domains in *NSP1* and *NUP116* mRNAs (**Fig. 1C**, grey bars). In an *in vitro* binding assay, synthetic biotinylated RNA probes encompassing these Hek2-binding sequences were further found to specifically pull down recombinant, purified Hek2 (**Fig. 1E**), but not a control protein (**Fig. S1E**).

Altogether, our data establish that a direct association with the hnRNP Hek2 specifically defines a subset of *NPC* mRNPs. Notably, the five Hek2-bound *NPC* mRNAs are coding for FG-Nups, which are critical for nucleocytoplasmic transport (Strawn et al., 2004).

### Hek2-dependent translational repression of *NPC* mRNAs and protein turnover define nucleoporins levels

We further investigated how Hek2 binding impacts on the fate of these particular *NPC* mRNAs. While previous studies have revealed that Hek2 associates with an important fraction of the transcriptome, the consequences of this recruitment for mRNA metabolism has only been documented in a few situations where Hek2 binding can cause increased mRNA stability (Hasegawa et al., 2008), asymmetrical localization (Irie et al., 2002) or translational repression (Paquin et al., 2007; Wolf et al., 2010).

To determine whether Hek2 binding influences the steady-state levels of *NPC* mRNAs, we first profiled the transcriptome of *hek2Δ* mutant yeast cells (**Fig. 2A**). Genome-wide, Hek2-bound mRNAs showed a tendency to be less abundant upon Hek2 inactivation (**Fig. S2A**), a trend not observed for Nab2-associated transcripts (**Fig. S2B**), highlighting the sensitivity and the specificity or our analysis. However, *NPC* mRNAs levels were not significantly affected by the absence of Hek2, whether or not they associate with this factor (**Fig. 2A**). We then compared the localization of *NPC* mRNAs in *wt* and *hek2Δ* cells using single molecule fluorescence *in situ* hybridization (**Fig. 2B**). Detection of *NSP1*, *NUP100* and *NUP133* mRNAs using specific sets of probes revealed a punctuate, cytoplasmic localization for these Nups-encoding transcripts in *wt* cells (**Fig. 2B**, top panels). Upon *HEK2* deletion, this random distribution, as well as the total number of detected RNA dots, were unchanged for both Hek2-bound (*NSP1, NUP100*) and unbound (*NUP133*) mRNAs (**Fig. 2B**, bottom panels). This set of data therefore establishes that Hek2 binding modulates neither the levels nor the localization of *NPC* mRNAs.

We then monitored the possible influence of Hek2 on *NPC* mRNAs translation using polysome fractionation on sucrose gradients, which resolve free mRNPs and ribosomal subunits from translation-engaged mRNAs (**Fig. 2C**, **Fig. S2C**). RT-qPCR analysis of the fractions of the *wt* polysome gradient revealed a bimodal distribution for Hek2-bound (**Fig. 2D**, **Fig. S2D,** black lines) and Hek2–unbound (**Fig. 2E**, **Fig. S2E**, black lines) *NPC* mRNAs. The largest fraction of *NPC* mRNAs migrated in the lightest fractions (*#1-6*), corresponding to free, untranslated mRNPs and resembling the pattern observed for the repressed *ASH1* mRNA (**Fig. 2D**). A less abundant fraction of *NPC* mRNAs peaked with polysomes-containing fractions (*#9-13*), similar to the peak of the well-translated *ACT1* mRNA (**Fig. 2E**). Puromycin-mediated dissociation of polysomes led to the disappearance of this second *NPC* mRNAs population, confirming that it corresponds to actively translated mRNPs (our unpublished data). Further analysis of the polysome profile from *hek2Δ* cells did not reveal any differences in the distribution of ribosomal species as compared to *wt* cells (**Fig. 2C**, **Fig. S2C**). Strikingly, *HEK2* inactivation decreased the amounts of translationally-repressed Hek2-bound *NPC* mRNAs (**Fig. 2D**, grey arrows) and triggered their redistribution in the translated population, with a peak in heavy polysomes fractions (≥4 ribosomes/mRNA; **Fig. 2D,** red arrows). This behavior was similar to the one reported for the Hek2-repressed *ASH1* mRNA (Paquin et al., 2007; see also **Fig. 2D**) and was not observed for mRNAs which are not bound by Hek2 (e.g. *NUP133* and *ACT1*, **Fig. 2E**).

**Figure 2.**
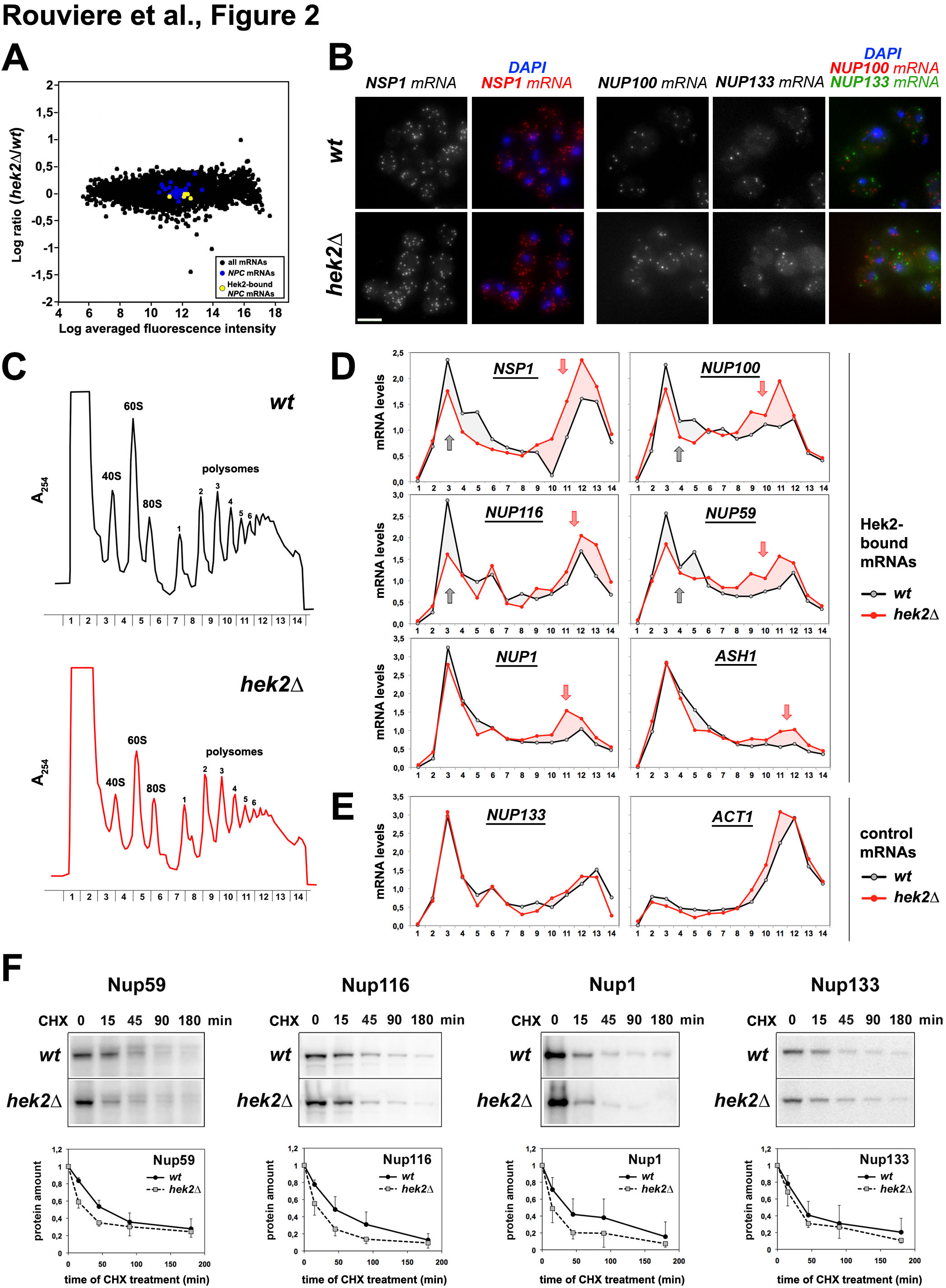
Hek2-dependent translational repression and protein turnover define nucleoporin levels. **A,** Transcriptome analysis of the *hek2Δ* mutant (MA plot). The Y-axis (M) is the averaged log2 of the *hek2Δ/wt* ratios calculated from two independent microarray hybridizations. The X-axis (A) is the log2 of the averaged fluorescence intensities. mRNAs encoding NPCs components are highlighted in two different colors depending on their association to Hek2 (from **Fig. 1**). **B**, Single molecule FISH was performed on *wt* and *hek2Δ* cells using set of probes specific for *NSP1*, *NUP100* or *NUP133* mRNAs. *NSP1* and *NUP100* probes were coupled to the Quasar570 fluorophore (red), while *NUP133* probes were coupled to Quasar670 (far red). z-projections are displayed, together with merged images with a nuclear staining (DAPI). Scale bar, 5 *μ*m. **C,** Polysome fractionation from *wt* and *hek2Δ* cells from the W303 background. The absorbance at 254nm (A_254_) recorded during the collection of the different fractions of the sucrose gradient is displayed. The positions of 40S, 60S, 80S ribosomal species and polysomes are indicated, as well as the number of ribosomes per mRNA in the polysomes fractions. **D**, Relative distribution of the *NSP1*, *NUP100*, *NUP116*, *NUP59*, *NUP1* and *ASH1* mRNAs in polysome gradients from the same *wt* (black lines) and *hekΔ* (red lines) cells. mRNAs amounts in each fraction were quantified by RT-qPCR, normalized to the sum of the fractions and to the distribution of a control spike RNA. Grey arrows indicate a decrease in the amounts of mRNAs found in the light fractions in *hek2Δ* cells. Red arrows point to an increase in the quantity of mRNAs found in the polysomes fractions of the mutant. These results are representative of four independent experiments (two performed in the W303 background, two in the BY4742 background; see **Fig. S2**). **E,** Same as **D,** for *NUP133* and *ACT1* control mRNAs. **F,** Protein levels of the indicated nucleoporins (Nup116, Nup1, Nup133) and of a GFP-tagged version of Nup59 were scored in *wt* and *hek2Δ* cells treated with cycloheximide (CHX) for the indicated time (minutes). *Top*, Whole cell extracts were analyzed by western blotting using anti-GLFG, anti-FSFG, anti-Nup133 or anti-GFP antibodies. *Bottom*, The relative amounts of the indicated proteins (mean ± SD; n=3) were quantified over the time following CHX treatment and are expressed relative to t=0. **See also Fig. S2.**

Having established that Hek2 binding onto its *NPC* target mRNAs contributes to their maintenance in a translationally-repressed state, we wondered whether it would affect the raw levels of their cognate nucleoporins. Notably, *HEK2* inactivation, while increasing the fraction of translated *NUP59*, *NUP116*, or *NUP1* mRNAs (**Fig. 2D**), did not trigger any drastic changes in the steady-state levels of the corresponding nucleoporins (see t=0 on **Fig. 2F;** our unpublished data). Since excess synthesis of subunits of multiprotein complexes can be buffered by increased protein turnover (Dephoure et al., 2014), we monitored the half-lives of these nucleoporins in *wt* and *hek2Δ* cells. Strikingly, the degradation rates of the three nucleoporins, as estimated from cycloheximide chase experiments, were higher in the absence of Hek2 (**Fig. 2F**), revealing that the enhanced synthesis of nucleoporins is attenuated by their increased turnover in these mutant cells. Consistently, the kinetics of degradation of Nup133, whose translation is independent from Hek2 activity, was unaffected in *hek2Δ* cells (**Fig. 2F**). The raw levels of this subset of nucleoporins are thereby tightly controlled by both Hek2-mediated translational control and protein degradation.

### Hek2 binding to *NPC* mRNAs requires prior desumoylation by the NPC-associated SUMO protease Ulp1

To determine whether Hek2 could transduce physiological signals that would modulate *NPC* mRNAs expression, we then looked for possible regulations of Hek2 function. Yck1-mediated phosphorylation of Hek2 was previously reported to disrupt its association with the *ASH1* mRNA at the bud cortex where this asymmetrically localized mRNA is targeted (Paquin et al., 2007). However, this plasma membrane-anchored kinase is unlikely to similarly target cytoplasm-localized *NPC* mRNPs (**Fig. 2B**). In view of the functional relationships between sumoylation and NPCs (Palancade and Doye, 2008) and of the multiple examples of nucleic acid-binding proteins whose activity is controlled by SUMO (Rouviere et al., 2013), we rather wondered whether Hek2 could be regulated by this modification.

To answer this question, cellular SUMO-conjugates were purified by denaturing Ni^2+^ chromatography from strains expressing a poly-histidine-tagged version of SUMO and a HA-tagged version of Hek2 (**Fig. 3A**). This assay specifically detected slower-migrating, mono-sumoylated versions of Hek2 in the SUMO-conjugates fraction of cells co-expressing Hek2-HA and His-SUMO (**Fig. 3B**). Strikingly, these sumoylated forms accumulated in *ulp1* mutant cells (**Fig. 3C**), demonstrating that this SUMO protease enables SUMO deconjugation from Hek2. To further determine the lysine residues that are modified by SUMO within this protein, we generated several plasmid-based *hek2* mutants where multiple lysines were mutated to arginines to prevent SUMO conjugation without disturbing the charge of the protein (**Fig. S3A**), and expressed them in *hek2Δ* cells. While mutations of all Hek2 lysines (*K1-30R*) completely abolished sumoylation, mutations of residues 19 to 30 (*K19-30R*), 25 to 30 (*K25-30R*) or 29/30 (*K29-30R*) were found to prevent the formation of most of the lower sumoylated version of Hek2 (**Fig. S3B**, lanes 5, 15, 32, 35), and mutations of lysines 8 to 18 (*K8-18R*), 13 to 18 (*K13-18R*) or 15 alone (*K15R*) strongly decreased its major upper sumoylation band (**Fig. S3B**, lanes 4, 13, 22, 24). Consistently, the *K15R K29-30R* combined mutant strongly reduced Hek2 sumoylation (**Fig. 3D**). Importantly, the turnover of Hek2 was unaffected in conditions where its sumoylation was enhanced (*ulp1* cells) or decreased (*hek2-K15 K29-30R* cells), demonstrating that this modification does not regulate its stability (**Fig. S3C**). Furthermore, Hek2 subcellular localization was unchanged in conditions of hypersumoylation (*ulp1* mutant, our unpublished data).

**Figure 3.**
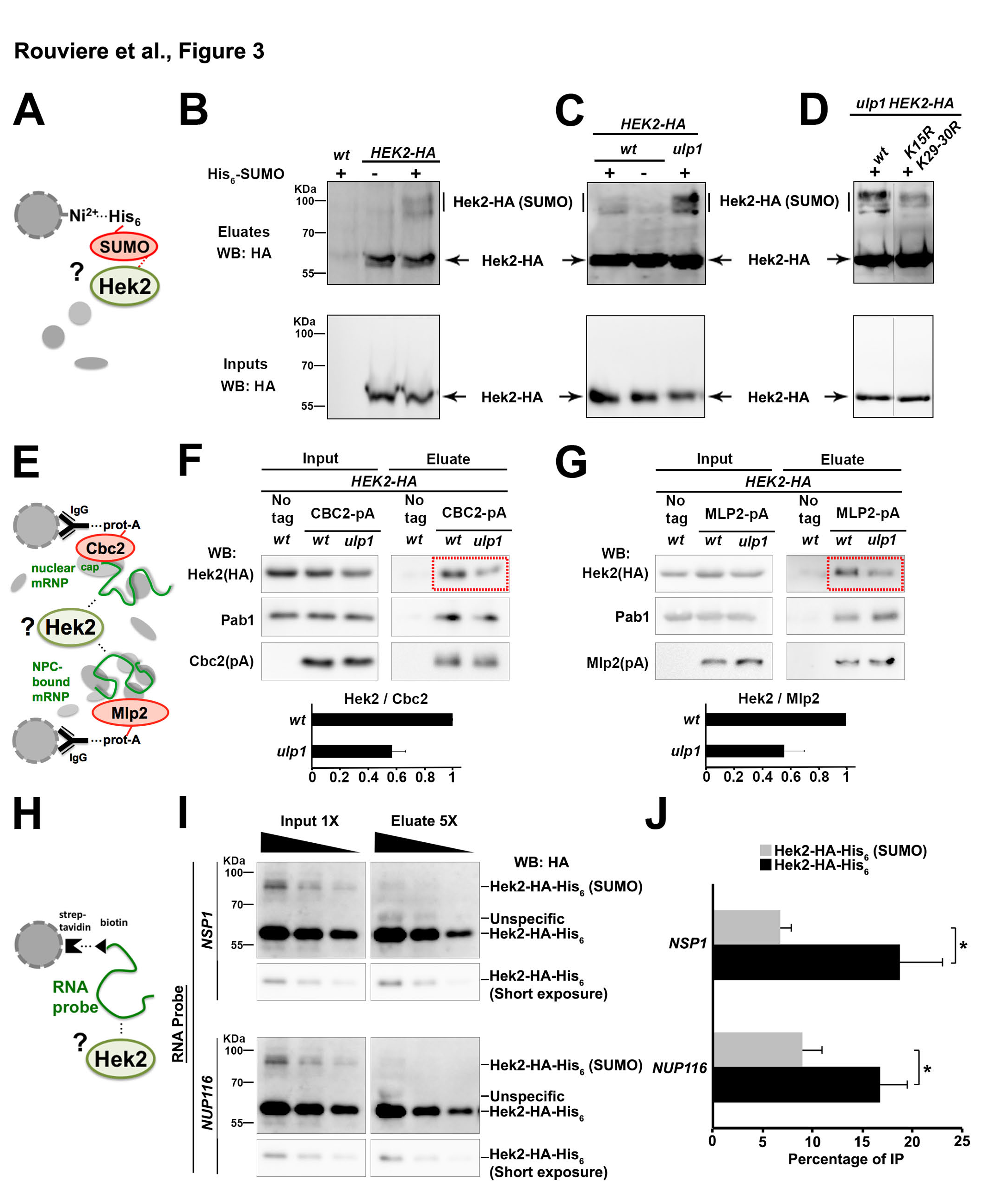
– Hek2 sumoylation prevents its association to mRNAs. **A,** Principle of the purification of sumoylated Hek2. Extracts from cells expressing a His-tagged version of SUMO were used for denaturing nickel chromatography**. B-D,** Extracts from *wt* and *HEK2-HA* cells (**B**), *HEK2-HA* and *HEK2-HA ulp1* cells (**C**), or *HEK2-HA ulp1* and *HEK2 K15R K29-30R-HA ulp1* cells (**D**) expressing or not His_6_-SUMO (+/-) were used for nickel chromatography. Total lysates (”Inputs”) and purified His-SUMO conjugates (”Eluates”) were analyzed by western blotting using anti-HA antibodies. The positions of the sumoylated and unmodified versions of Hek2-HA, as well as molecular weights, are indicated. The apparent molecular weights of the modified versions of Hek2 are compatible with mono-sumoylations occurring on distinct residues. Note the non-specific binding of a fraction of non-sumoylated Hek2-HA (also observed in the absence of His-SUMO, 2^nd^ lanes in panels **B** and **C**), a classical issue in SUMO-conjugates purification. *ulp1* mutant cells carry the *ulp1-333* thermosensitive allele that disturbs both Ulp1 activity and NPC localization. **E,** Principle of the mRNP purification procedure. Cbc2 or Mlp2 are purified through a protein-A tag, and the protein content of the associated mRNPs is analyzed by western blot. Note that RNAse A treatment experiments confirmed the RNA dependence of the interactions scored in such assays (Bretes et al., 2014). **F-G,** *Top*, Soluble extracts (”Inputs”, left panels) and Cbc2-pA-associated mRNPs (**F**) or Mlp2-pA-associated mRNPs (**G**) (”Eluate”, right panels) isolated from *wt* and *ulp1* cells were analyzed by immunoblotting using the indicated antibodies. *Bottom*, The relative amounts of Hek2 associated to Cbc2-and Mlp2-bound mRNPs (mean ± SD; n=3 for Cbc2-pA; n=2 for Mlp2-pA; set to 1 for *wt*) are represented. **H,** Principle of the *in vitro* RNA binding assay. **I**, An *in vitro* sumoylation mixture containing both unmodified and sumoylated Hek2 was incubated with streptavidin beads previously coated with biotinylated RNA probes encompassing Hek2-binding sites from *NSP1* or *NUP116*. Decreasing amounts of input and eluate fractions were loaded to allow quantification. **J**, Percentages of IP (mean ± SD; n=3) are the ratios between unmodified (or sumoylated) Hek2 amounts in the eluate and in the input fractions. * P&#x003C;0,05 (Welch’s t-test). **See also Fig. S3**.

In order to determine whether Hek2 sumoylation could rather regulate its interaction with its target mRNAs, we combined the following approaches. First, we purified two different subsets of mRNPs from *wt* and *ulp1* cells and analyzed their association with Hek2 (**Fig. 3E**). mRNPs were isolated using as baits either Cbc2, a subunit of the nuclear cap-binding complex (Cbc2-pA, **Fig. 3F**), or Mlp2, which anchors mRNPs to NPCs prior to nuclear export (Mlp2-pA, **Fig. 3G**; Bretes et al., 2014). Strikingly, *ULP1* loss-of-function triggered a clear decrease in the amounts of Hek2 recovered in both complexes (**Fig. 3F-G**), while it did not affect the recruitment of canonical mRNP components such as the poly-A binding protein Pab1, in agreement with our previous study (Bretes et al., 2014). Second, we specifically looked at the association of Hek2 with *NPC* mRNAs in *wt* and *ulp1* cells through Hek2-pA immunoprecipitation followed by RT-qPCR. This assay further confirmed that *ULP1* inactivation leads to a decrease in the association of Hek2 with its target mRNAs (**Fig. S3D**).

These two experiments demonstrate that the SUMO protease Ulp1 is required for both Hek2 desumoylation and binding to *NPC* mRNAs, suggesting that this association could be directly repressed by SUMO. To further challenge this hypothesis, we went on to compare the binding of unmodified and sumoylated Hek2 to *NPC* mRNAs in a reconstituted *in vitro* assay (**Fig. 3H**). For this purpose, we first achieved the *in vitro* sumoylation of recombinant Hek2 in the presence of purified versions of the SUMO-activating enzyme (Aos1-Uba2), the SUMO-conjugating enzyme (Ubc9) and SUMO, partly reproducing the observed *in vivo* sumoylation pattern (**Fig. S3E**, first lane). When further used in the *in vitro* RNA binding assay, the sumoylated version of Hek2 was unambiguously less prone to bind RNA that its unmodified counterpart (**Fig. 3I-J**). Altogether, our data thereby establish that Hek2 sumoylation negatively regulates its association to *NPC* mRNAs and that Ulp1 desumoylating activity is required for optimal binding.

### Compromised NPC integrity alters the levels of the SUMO protease Ulp1 and of active Hek2

The fact that the SUMO protease that controls the binding of Hek2 to *NPC* mRNAs is itself associated to nuclear pores prompted us to test whether it could be part of a feedback mechanism sensing NPC integrity and further modulating Nups biogenesis. We therefore asked whether mutant or physiological situations associated with defects in nuclear pore functions would result in changes in the activity of Ulp1 towards Hek2.

Mutants of distinct NPC subcomplexes, e.g. the outer ring Nup84 complex and the nuclear basket Nup60-Mlp1/2 complex, were previously shown to exhibit decreased levels of Ulp1 at the nuclear envelope (Zhao et al., 2004; Palancade et al., 2007). To complement these findings, we systematically analyzed the localization of Ulp1 in *ΔFG* mutants in which the genetic removal of FG domains from specific nucleoporins leads to defects in nucleocytoplasmic transport, including karyopherin-dependent import (Strawn et al., 2004). In *wt* cells, the GFP-tagged version of Ulp1 exhibited a discontinuous rim-like staining of the nuclear periphery typical of its NPC-associated localization (**Fig. 4A**). In most *ΔFG* mutants however, the Ulp1-GFP nuclear envelope staining was significantly reduced (**Fig. 4A-B**). This phenotype was unlikely to be caused by a reduction in the number of NPCs, according to a previous characterization of these mutants (Strawn et al., 2004), but rather reflected a decrease in the karyopherin-dependent import step that precedes Ulp1 anchoring at NPCs. Consistently, we did not observed this reduced Ulp1 staining in the *nup1ΔFG* mutant (**Fig. 4A-B**) which is unexpected to impair karyopherin function (Strawn et al., 2004).

**Figure 4.**
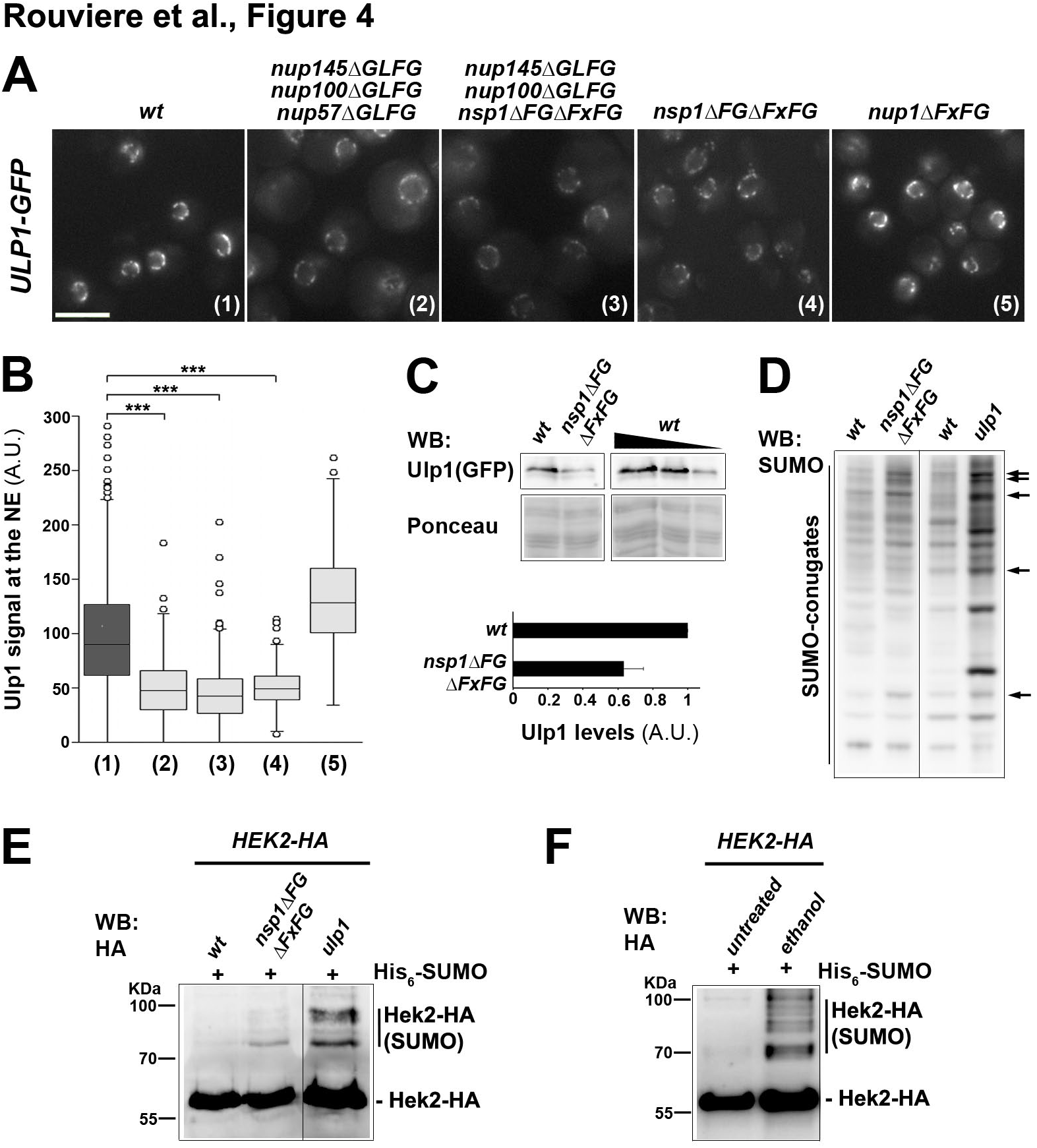
– Defects in nuclear pore integrity impact Ulp1 activity and Hek2 sumoylation. **A,** Fluorescence microscopy analysis of Ulp1-GFP in *wt*, *nup145ΔGLFG nup100ΔGLFG nup57ΔGLFG*, *nup145ΔGLFG nup100ΔGLFG nsp1ΔFGΔFxFG*, *nsp1ΔFG·FxFG* and *nup1ΔFxFG* cells grown at 30°C. Scale bar, 5 *μ*m. **B,** Quantification of the Ulp1 nuclear envelope fluorescence intensity in the different strains. The numbers refer to the genotypes as depicted in **A**. For each strain, at least 150 cells were analyzed. *** P&#x003C;0,001 (Mann Whitney Wilcoxon test). **C**, Ulp1-GFP amounts were measured in *wt* and *nsp1ΔFGΔFxFG* cells by western blotting using anti-GFP antibody (*top panel*). Ponceau staining was used as a loading control (*lower panel*). A serial dilution of the *wt* sample was used for quantification. Ulp1-GFP amounts normalized to ponceau (mean ± SD; n=2; set to 1 for *wt*) are represented. **D,** Whole cell extracts of the indicated strains were analyzed by western blotting using anti-SUMO antibodies. The bands that are modified in the *nsp1ΔFGΔFxFG* mutant are also typically altered in *ulp1* cells (arrows). **E**, Hek2 sumoylation was detected in *wt* and *nsp1ΔFGΔFxFG* cells as in **Fig. 3**. The pattern of Hek2 sumoylation in *ulp1* cells was analyzed as a control. The positions of the sumoylated and unmodified versions of Hek2-HA, as well as molecular weights, are indicated. **F,** Hek2 sumoylation was detected in *wt* cells, either untreated, or treated with 10% ethanol for 1 hour.

To further characterize this phenotype, we pursued the analysis of the *nsp1ΔFGΔFxFG* mutant in which removal of the FG domains from a single nucleoporin is sufficient to decrease Ulp1 levels at the nuclear envelope (**Fig. 4A-B**). In agreement with the previous reported interdependence between Ulp1 NPC localization and stability (Zhao et al., 2004; Palancade et al., 2007), western-blot analysis of this *nsp1ΔFGΔFxFG* mutant further revealed a reduction in the total amounts of cellular Ulp1 as compared to *wt* cells (**Fig. 4C**). Consistently, analysis of the global pattern of cellular SUMO conjugation in this same mutant highlighted a number of discrete changes, in particular the accumulation of high-molecular weight SUMO conjugates, resembling those caused by *ULP1* inactivation (**Fig. 4D,** arrows). We then wondered whether the changes in Ulp1 levels and activity detected in this mutant were sufficient to modulate Hek2 sumoylation. Remarkably, *nsp1ΔFGΔFxFG* cells exhibited a clear increase in the levels of sumoylated Hek2 (**Fig. 4E**). Loss of NPC integrity upon genetic alteration of several distinct NPC components can therefore impact the levels of active Ulp1, which is sufficient to trigger the accumulation of sumoylated, inactive versions of Hek2.

We finally asked whether physiological changes in NPC integrity could also lead to the accumulation of inactive Hek2 in *wt* cells. Environmental stresses can trigger changes in NPC integrity, as exemplified by the specific delocalization of certain NPC components, including Ulp1, upon exposition to elevated alcohol levels (Izawa et al., 2004; Takemura et al., 2004; Sydorskyy et al., 2010). We then analyzed the sumoylation levels of Hek2 in *wt* cells exposed to ethanol stress (**Fig. 4F**). Strikingly, increased levels of sumoylated Hek2 were detected in this situation (**Fig. 4F**). Changes in NPC integrity, triggered by either genetic alterations or physiological changes, can thereby translate into the accumulation of inactive versions of Hek2.

## DISCUSSION

By combining the analysis of genomic data with *in vivo* and *in vitro* interaction assays, we have established that a subset of the mRNAs that encode the subunits of nuclear pores display a unique mRNP composition characterized by the binding of the hnRNP Hek2/Khd1 (**Fig. 1**). This conserved RNA-binding protein was previously reported to have various effects on the metabolism of its target mRNAs (Irie et al., 2002; Paquin et al., 2007; Hasegawa et al., 2008; Wolf et al., 2010), possibly reflecting coregulations involving other RBPs (Ito et al., 2011), including the Hek2 paralogue Pbp2/Hek1, or transcript specificities, as in the case of the bud-localized mRNA *ASH1*. Here, we show that Hek2 binding to Nup-encoding mRNAs affects neither their steady-state levels nor their subcellular localization (**Fig. 2A-B**), in contrast with other target mRNAs (**Fig. S2A-B**; Irie et al., 2002). In contrast, Hek2 binding appears to regulate the translation of *NPC* mRNAs. Indeed, upon *HEK2* inactivation, the percentage of translated Hek2 target mRNAs increases and peaks with the heavy polysomes containing the most actively translating ribosomes, a phenotype that is not observed for control transcripts (**Fig. 2D-E**). In this frame, the regulation of *NPC* mRNAs is reminiscent of the one scored for *ASH1* and *FLO11*, two mRNAs for which Hek2 binding represses translation initiation (**Fig. 2D**; Paquin et al., 2007; Wolf et al., 2010). Moreover, our study uncovers that in *wt* cells, *NPC* mRNAs distribute in two populations, one being actively translated and the other translationally-repressed. Such a bimodal distribution is rather uncommon in yeast, in which whole-genome polysomal profiles previously revealed that most mRNAs are associated with translating ribosomes during exponential growth (Arava et al., 2003), and likely indicates undergoing translational controls. However, it has to be noted that Hek2 binding is unlikely to be the only determinant of this particular translational regulation. Indeed, a large fraction of each Hek2-bound mRNAs (e.g. *NSP1* and *NUP1*, **Fig. 2D**) remains untranslated in the absence of Hek2. In addition, the *NPC* mRNAs that are not among Hek2 preferred targets (e.g. *NUP133*, **Fig. 2E**) also exist for the most part in a translation-inactive fraction. Whether alternate RBPs, specific for distinct subsets of *NPC* mRNAs, or other layers of regulations also partake in the fine-tuning of the translation of these transcripts remains to be investigated.

While Hek2 represses *NPC* mRNAs translation, protein turnover also contributes to the definition of the cellular levels of nucleoporins. Indeed, excess Nups likely synthesized in the absence of Hek2-dependent translational repression appear to be buffered by an increase in their degradation rates (**Fig. 2F**). This mechanism is reminiscent of the post-translational attenuation described to occur for multiprotein complex subunits when they are naturally produced in super-stoichiometric amounts (McShane et al., 2016), or overexpressed due to genomic amplification (Dephoure et al., 2014). Excess subunits of NPCs, which do not assemble into stable complexes and could be possibly unfolded, are thereby expected to undergo increased ubiquitin-dependent, proteasome-mediated degradation. Several conserved ubiquitin ligases are susceptible to partake in this process, including (i) Hul5 and San1, which recognize misfolded proteins in the cytoplasm and the nucleus, respectively (Fang et al., 2011; Rosenbaum et al., 2011); (ii) Tom1, which couples ubiquitin to unassembled ribosomal proteins (Sung et al., 2016) or (iii) any yet-to-be characterized quality control factor specialized in the degradation of orphan polypeptides, as recently identified in mammals (Hampton and Dargemont, 2017). The fact that the cellular concentration of Hek2-regulated nucleoporins such as Nup59, Nup1 and Nup116 is tightly restricted by both translational repression and protein degradation suggests that their accumulation could be detrimental, either because they could interfere with the assembly of NPCs, or because these hydrophobic proteins would be prone to the formation of toxic aggregates. Possibly in line with both hypotheses, overproduction of Nup170, a direct partner of Nup59 at the NPC, was reported to trigger the accumulation of cytoplasmic foci containing distinct unassembled nucleoporins (Flemming et al., 2009), while overexpressed Nup59 was found to accumulate within cytoplasmic structures (Marelli et al., 2001).

In agreement with the physiological importance of such Hek2-mediated regulations, it is not surprising that the activity of this protein is itself under control. We found that sumoylation of Hek2 occurs on two different domains, thus generating two distinct monosumoylated versions of the protein (**Fig. 3B-D**, **Fig. S3A-B**). Both modified regions are located at the vicinity of the third K-homology (KH) domain (**Fig. S3A**), the major RNA-interacting motif of the protein (Hasegawa et al., 2008), providing a possible molecular rationale for the SUMO-mediated decrease in RNA binding scored *in vivo* (**Fig. 3F-G**, **Fig. S3D**) and *in vitro* (**Fig. 3I-J**). In this respect, inhibition of RNA recognition could be caused by steric hindrance, as already reported for several sumoylated DNA-or RNA-binding proteins (Rouviere et al., 2013), or, alternatively, occur through changes in the oligomerization status of the protein, as proposed in the case of human hnRNP C1 (Vassileva and Matunis, 2004). Furthermore, the spatio-temporal control of Hek2 function is likely to depend on a combination of post-translational modifications including, besides its sumoylation, its reported phosphorylation by Yck1 (Paquin et al., 2007) and its ubiquitination detected in proteome-wide analyses (Starita et al., 2012). Finally, in view of the compartmentalization of the enzymes of the SUMO pathway, Hek2 sumoylation is likely to occur mainly inside the nucleus, while its desumoylation by Ulp1 would favor its binding onto mRNAs at the nucleoplasmic side of NPCs (**Fig. S4A**). The cytoplasmic fate of certain mRNPs would then be determined prior to export, as in the case of *ASH1* whose asymmetrical localization and translation depends on Hek2 binding. This molecular mechanism could also explain why *ASH1* asymmetry requires Nup60 (Powrie et al., 2011), since this nucleoporin is one of the major determinants of Ulp1 stability at NPCs (Zhao et al., 2004; Palancade et al., 2007).

The control of Hek2 function through Ulp1-mediated desumoylation is also likely to adjust its RNA binding activity in response to the status of nuclear pores in the cell. Since several distinct nucleoporin subcomplexes are indeed required to position and stabilize Ulp1 at the pore (Zhao et al., 2004; Palancade et al., 2007; **Fig. 4A-B**), the level of activity of this SUMO protease provides a read out for the number and the functionality of NPCs. Consistently, changes in NPC composition in mutant or perturbed physiological situations impact Ulp1 activity and trigger the accumulation of sumoylated, inactive versions of Hek2 (**Fig. 4E-F**). In view of the function of Hek2 in controlling NPC mRNAs translation (**Fig. 2**), this could in turn result in the increased synthesis of nucleoporins in a feedback process (**Fig. S4B**). Their recruitment into NPCs would then compete their proteasomal degradation and contribute to restore NPC integrity. Strikingly, some of the nucleoporins that are targeted by this mechanism appear to be the most limiting ones for completing fully-assembled NPCs (**Fig. S4C**). Among them, Nsp1 is also critical to define NPC number during the asymmetric division of budding yeast (Colombi et al., 2013; Makio et al., 2013). While the pathway described here could indeed connect the cellular availability of specific nucleoporins to the status of NPCs, other quality control mechanisms are known to control NPC homeostasis. In yeast, aberrant NPC assembly intermediates are cleared from the nuclear envelope by the activity of ESCRT-III/Vps4 complexes (Webster et al., 2014), while in mammals, defects in the assembly of nuclear pore baskets triggers a cell cycle delay (Mackay et al., 2010).

Localization of SUMO proteases at NPCs has been conserved in all eukaryotes (Palancade and Doye, 2008) and also involves several distinct NPC-associated determinants in mammalian cells (Goeres et al., 2011; Hang and Dasso, 2002). Sumoylation of KH-domain containing Hek2 orthologues such as hnRNP K, hnRNP E1 and hnRNP E2 has also been reported (Bruderer et al., 2011; Li et al., 2004; Pelisch et al., 2012). Strikingly, hnRNP K desumoylation involves SENP2, the NPC-localized orthologue of Ulp1 in mammals (Lee et al., 2012). In view of the association between hnRNP K and a subset of *NPC* mRNAs in a genome-wide survey of human RBPs (Van Nostrand et al., 2017), the conservation of the pathway described here will certainly deserve further investigation.

## AUTHOR CONTRIBUTIONS

Conceptualization, J.R., B.P.; Methodology, J.R., B.C., B.P.; Investigation, J.R., M.B., B.C, F.D., B.P.; Formal analysis, J.R., A.T., F.D., B.P.; Writing, J.R., B.P.; Visualization, J.R., B.P.; Funding acquisition, B.P.; Supervision, B.C., B.P.

## ACKNOWLEDGMENTS

We thank P. Chartrand, C. Dargemont, V. Doye, E. Johnson and S. Wente for reagents; L. Fusée and G. Lelandais for help with *in vitro* experiments and data normalization, respectively; D. Tollervey for CRAC analyses; A. Babour, V. Doye, T.H. Jensen and R. Rothstein for discussion and critical reading of the manuscript. This work was supported by: CNRS (to B.P.); Fondation ARC pour la Recherche sur le Cancer (to B.P.); Ligue Nationale contre le Cancer (to B.P.; fellowship to J.R.); Ecole Doctorale “Structure et Dynamique des Systèmes Vivants” (#576), Université Paris-Sud, Université Paris-Saclay (fellowship to J.R.); and the ‘Who am I?’ laboratory of excellence (grant numbers ANR-11-LABX-0071, ANR-11-IDEX-0005-02, fellowship to J.R).

## METHODS

### Yeast strains and plasmids

Unless otherwise indicated, all the strains used in this study (listed in **Table S1**) are isogenic to BY4742/BY4741 and were grown in standard culture conditions. When indicated, cycloheximide (0.1mg/mL, Sigma) or ethanol (10% v/v) were added to the medium for the indicated time. Construction of plasmids (listed in **Table S2**) was performed using standard PCR-based molecular cloning techniques.

### Bioinformatic analysis of RNA immunoprecipitation datasets

RNA immunoprecipitation (RIP), cross-linking immunoprecipitation (CLIP) or crosslinking and analysis of cDNA (CRAC) data were collected for the following RNA-binding proteins: Yra1 (RIP followed by microarray analysis, one replicate; Hieronymus and Silver, 2003), Nab2 (CRAC, three replicates, Tuck and Tollervey, 2013), Npl3 (RIP followed by microarray analysis, one replicate, Kim Guisbert et al., 2005), Nab4/Hrp1 (RIP followed by microarray analysis, one replicate, Kim Guisbert et al., 2005), Mex67 (CRAC, three replicates, Tuck and Tollervey, 2013), Sto1 (CRAC, three replicates, Tuck and Tollervey, 2013), Xrn1 (CRAC, two replicates, Tuck and Tollervey, 2013), Ski2 (CRAC, four replicates, Tuck and Tollervey, 2013), Mtr4 (CRAC, three replicates, Tuck and Tollervey, 2013), Hek2 (CRAC, one replicate, Tuck and Tollervey, 2013; CLIP, one replicate, Wolf et al., 2010; RIP followed by microarray analysis, one replicate in two distinct studies, Hasegawa et al., 2008, Hogan et al., 2008). For each dataset, all protein-coding RNAs were ranked and given a color according to their relative binding to the corresponding RBP. Scores available from microarray or sequencing analyses (Hasegawa et al., 2008; Hieronymus and Silver, 2003; Hogan et al., 2008; Kim Guisbert et al., 2005) were used to split the RNAs in four equally sized groups corresponding respectively to “high” (light yellow), “medium” (dark yellow), “low” (dark blue) and “very low/no” (light blue) binding. CLIP data were used to define bound (light yellow) and unbound (light blue) mRNAs according to the published peak calling analysis (Wolf et al., 2010). CRAC hits were first normalized by hits per million within each RBP CRAC dataset, then for each mRNA (Σi^2^=1) to account for differences in mRNA abundances, and scaled to occupy the 0-1 range. Colors ranging from light blue (0) to light yellow (1) were used to depict the binding of a given mRNA to a RBP. Binding categories were further displayed for *NPC* mRNAs (**Fig. 1A**) or proteasome/exosome RNAs (**Fig. S1C**). Gene set enrichment analyses were performed as described (Subramanian et al., 2005). The MEME software (v4.11.3; Bailey et al., 2009) was applied to the sequences of *NUP59, NUP116, NUP1, NSP1* and *NUP100* mRNAs. Out of 6 retrieved motifs, 5 corresponded to FG-coding sequences while one, found with an e-value of 4.8e-7, matched the known Hek2-binding site (**Fig. 1D**).

### mRNP and RNA immunoprecipitation

Cbc2-pA-and Mlp2-pA-associated mRNPs complexes were purified as previously described (Bretes et al., 2014) except that extracts were incubated with IgG-conjugated magnetic beads for 10 instead of 30 min. Hpr1 RNA immunoprecipitation was performed as previously described (Bretes et al., 2014). Hek2-pA associated mRNA purifications were performed according to the same procedure in the presence of RNAsin (Promega, 40U per mL of buffer). Total and immunoprecipitated RNAs were purified with the Nucleospin RNAII kit (Macherey Nagel) and reverse-transcribed with Superscript II reverse transcriptase (Life Technologies). cDNA were further quantified by real-time PCR with a LightCycler 480 system (Roche) according to the manufacturer’s instructions. The sequences of the primers used for qPCR in this study are listed in **Table S3**. Controls without reverse transcriptase allowed estimating the lack of contaminating DNA.

### Polysome profiling analysis

The protocol was adapted from published procedures (Kuhn et al., 2001; Arava et al., 2003). 100 ml cultures were grown in YPD media to midlog phase (OD_600_ = 0.4-0.6). Prior to harvest, cycloheximide (CHX) (Sigma) was added to final a concentration of 0.1mg/ml. All subsequent procedures were carried out on ice with pre-chilled tubes and buffers. Cultures were cooled on ice and pelleted by centrifugation at 2,600*g* for 5 min at 4°C. Pellets were washed twice in 2.5 ml of ice cold lysis buffer (Tris-HCl 20mM, KCl 140mM, MgCl_2_ 1.5mM, Triton X-100 1% [vol/vol], DTT 0.5mM, CHX 0.1mg/ml and Heparin 1mg/ml), resuspended in 0.7 ml of ice cold lysis buffer and lysed by bead beating using a Fastprep (Qbiogene, 3x30s). Cell debris and glass beads were removed by centrifugation at 2,600*g* for 5 min at 4°C. The supernatant was transferred to a 1.5 ml tube and clarified by centrifugation at 10,000*g* for 10 min at 4°C. 10 A_254_ units of extract were layered onto an 11ml 20-50% (wt/vol) sucrose gradient prepared in the lysis buffer without Triton X-100. The samples were ultra-centrifuged at 39,000*g* for 2.5h at 4°C in a SW41 rotor. The gradients were fractionated in 14 fractions of 0.9 ml using an ISCO fractionation system with concomitant measurement of A_254_. Total lysates and fractions were supplemented with 50 *μ*l of NH_4_Ac 3M, 5 ng of Luciferase RNA (Promega), 1 *μ*l of Glycoblue (Ambion) and 1.2 ml of Ethanol. Samples were vortexed and precipitated overnight at −20°C. The pellets were collected by centrifugation at 10,000*g* for 10 min at 4°C, washed once in 75% ethanol and resuspended in 100 *μ*l DEPC-treated H_2_O. RNAs were further purified using the Nucleospin RNAII kit (Macherey-Nagel) following the RNA clean-up procedure. Equal volumes of all samples were reverse transcribed with Superscript II reverse transcriptase (Life Technologies) and cDNA were further quantified by real-time PCR as described above.

### Recombinant protein production

His and GST fusion proteins were expressed in Rosetta (DE3) *E. coli* cells transformed with the corresponding plasmids and grown in LB medium supplemented with the required antibiotics. Expression of the recombinant proteins was achieved by submitting bacterial cultures to cold and chemical shocks (4°C, 2% ethanol), and inducing them with 0.2mM isopropyl-β-D-thiogalactopyranoside at 23°C for 4h. Bacterial pellets were collected by centrifugation and frozen in liquid nitrogen. Pellets were resuspended either in His buffer (Na_2_HPO4 20mM pH 7.5, NaCl 500mM, imidazole 10mM, triton X-100 0.2%, MgCl2 1mM, protease inhibitors cocktail, Roche) or GST buffer (Tris HCl 50mM pH7.5, Triton 0.1%, KCl 10mM, Glycerol 10%, DTT 1mM, protease inhibitors cocktail, Roche), treated with 0.5mg/mL lysosyme for 1h at 4°C and lysed by sonication. His-tagged proteins were further solubilized by adding 0.5% Sarkosyl for 15min at 4°C, followed by the addition of 0.8% Triton X-100. Lysates were cleared by centrifugation at 10,000*g* for 20min at 4°C. His-tagged proteins were purified on Ni-NTA agarose (Qiagen) for 2h at 4°C. Beads were then washed twice with His buffer and eluted four time with the same buffer containing 500mM imidazole and 1% Triton X-100. GST fusion proteins were purified in the presence of 550mM NaCl on Gluthation sepharose (GE Healthcare) for 1h30 at 4°C. Beads were then washed tree times with GST buffer containing 500mM NaCl, and eluted four time 15min in Tris HCl 50mM pH8, NaCl 500mM, Triton 0.1%, glycerol 10%, gluthation 15mM. Following purification, His and GST fusion proteins were dialysed over night at 4°C against Hepes KOH 20mM pH7.9, KCl 0.1M, DTT 0.1mM, and 10% glycerol was added before storage at −80°C.

### *In vitro* RNA binding assay

*In vitro* RNA binding assays were performed according to a published procedure (Mehta and Driscoll, 1998). Streptavidin dynabeads (Invitrogen) were washed three times in 0.1M NaOH, 0.05M NaCl and once in 0.1M NaCl. 2*μ*g of biotinylated RNA (encompassing Hek2-binding sites on *NSP1* or *NUP116* mRNAs; Integrated DNA Technologies) were bound to 10*μ*L of beads in RNA binding buffer (Tris HCl 5mM pH 7.5, NaCl 1M, RNAsin 40U/mL) for 30min at room temperature. The conjugated beads were then washed four times in RNA binding buffer and incubated in protein binding buffer (Hepes 50mM pH7.5, NaCl 100mM, MgCl_2_ 1mM, glycerol 10%, DTT 0.5mM, PMSF 0.1mM, BSA 0.1%, RNAsin 40U/mL) for 15min at 4°C for saturation. Beads were then incubated in protein binding buffer containing 1mg/mL heparin and ∼2 pmol of recombinant Hek2 for 30min at 4°C. Beads were then washed five times with protein binding buffer containing 1mg/mL heparin and eluted in SDS sample buffer.

### Sumoylation assays

SUMO-conjugates were isolated from yeast cells expressing a His-tagged version of SUMO using nickel agarose denaturing chromatography as previously described (Bretes et al., 2014). *In vitro* sumoylation was performed as reported (Rothenbusch et al., 2012): briefly, 3*μ*g of recombinant Hek2 were mixed with 300nM of recombinant E1 enzyme (Aos1/Uba2), 700nM of recombinant E2 enzyme (Ubc9) and 10mM of a mutated version of Smt3 (K11,15,19R) less prone to form poly-SUMO chains, in the presence of 5mM ATP in a sumoylation buffer (BisTris 50mM pH 6.5, NaCl 100mM, MgCl_2_ 10mM, DTT 0.1mM). The reaction was then incubated for 3h at 37°C and either stopped by addition of SDS sample buffer, or further used for *in vitro* RNA binding assays.

### Protein extraction and Western blot analysis

Total protein extraction from yeast cells was performed by the NaOH–TCA lysis method (Ulrich and Davies, 2009). Samples were separated on 10% or 4–12% SDS-PAGE gels and transferred to nitrocellulose or PVDF membranes. Western-blot was performed using the following antibodies: polyclonal anti-GLFG (to detect Nup116; Grandi et al., 1995), 1:500; polyclonal anti-FSFG (to detect Nup1; Schlaich and Hurt, 1995), 1:4000; polyclonal anti-Nup133 (Belgareh and Doye, 1997), 1:500; monoclonal anti-Pab1 (1G1, Santa-Cruz), 1:1000 ; polyclonal anti-SUMO (Bonnet et al., 2015), 1:2000 ; monoclonal anti-HA (16B12, Covance), 1:1000 ; monoclonal anti-GFP (7.1 &#x0026; 13.1, Roche Diagnostics), 1:500 ; monoclonal anti-GST (4C10, Covance), 1:1000 ; rabbit IgG-HRP polyclonal antibody (to detect protein-A-tagged proteins, Dakocytomation), 1:5000. Quantification of signals was performed based on serial dilutions of reference samples using the ImageJ software.

### Gene expression analyses

Total RNAs were extracted from yeast cultures using Nucleospin RNA II (Macherey Nagel). Reverse transcription and cDNA quantification were performed as described above for RNA immunoprecipitation. Transcriptome analysis was achieved using microarrays as previously reported (Bretes et al., 2014). The *hek2Δ* versus *wt* comparison was performed twice using independent samples and dye swap. The averaged log_2_ of the mutant/wild-type ratios and the standard deviation between the two replicates were calculated for each gene. The genes showing a standard deviation > 0.5 were removed from the dataset. The complete microarray data are available in the ArrayExpress database (www.ebi.ac.uk/arrayexpress) under accession number E-MTAB-6065. Comparisons of *hek2Δ* transcriptome with Hek2 and Nab2 binding profiles were realized using published datasets (Hasegawa et al., 2008; Kim Guisbert et al., 2005). Transcripts were split in four equally sized groups corresponding respectively to “strong”, “medium”, “low” and “very low/non” binding. For each category, the log_2_ of the mutant/wild-type ratios of the different transcripts were represented as a box plot.

### Cell imaging

Single molecule fluorescence *in situ* hybridization (smFISH) was carried out on fixed cells using Stellaris Custom Probe Sets and RNA FISH buffers, according to the manufacturer’s instructions (Biosearch Technologies). For both smFISH and live imaging of the nuclear envelope, wide-field fluorescence images were acquired using a DM6000B Leica microscope with a 100X, NA 1.4 (HCX Plan-Apo) oil immersion objective and a CCD camera (CoolSNAP HQ; Photometrics). z-stacks sections of 0.2 *μ*m were acquired using a piezoelectric motor (LVDT; Physik Instrument) mounted underneath the objective lens. Images were scaled equivalently and 3D-projected using ImageJ, and further processed with Photoshop CS6 13.0 x64 software (Adobe). Nuclear envelope intensities were determined with ImageJ following subtraction of the cytoplasmic background.

### Statistics

(n) values correspond to the number of biological replicates (e.g. independent yeast cultures) and are indicated in the corresponding figure legends. Error bars correspond to standard deviations. The two-tailed Welch’s t-test, which allows unequal variance, was used to compare RNA binding efficiencies *in vitro* or *in vivo* (**Fig. 1E**; **Fig. 3J**; **Fig. S3D**). A Mann-Whitney-Wilcoxon test was used to compare Ulp1 nuclear envelope intensities in different strains (**Fig. 4B**) and RNA expression fold-changes upon *HEK2* deletion (**Fig. S2A-B**). Standard conventions for symbols indicating statistical significance were used: *P≤0.05; **P≤0.01; ***P≤0.001; N.S, not significant.

**Table S1:** Yeast strains used in this study

| Strain code | Name | Relevant genotype | Source |
| --- | --- | --- | --- |
| BY4742 / BY4741 | *wt* |  | Euroscarf |
| Y13058 | *hek2Δ* | *hek2::kanMX* | Euroscarf |
| JR154 | *HEK2-pA* | *hek2::kanMX*<br>*pRS316-HEK2-protA* | This study * |
| Y14072 | *hpr1Δ* | *hpr1::kanMX* | Euroscarf |
| K5552 | *wt* | **(W303)** *ASH1-MYC9* | (Paquin et al., 2007) |
| YV1862 | *hek2Δ* | **(W303)** *ASH1-MYC9 hek2::kanMX* | This study (a) |
| YV2083 | *NUP59-GFP* | *NUP59-GFP::HIS3MX* | Invitrogen |
| YV2084 | *NUP59-GFP hek2Δ* | *NUP59-GFP::HIS3MX hek2::kanMX* | This study * |
| YV1593 | *HEK2-3HA* | *HEK2-3HA::kanMX* | This study (b) |
| YV1262 | *ulp1* | *ulp1::kanMX*<br>*YCpLac111-ulp1-333 (LEU2)* | (Bretes et al., 2014) |
| YV1432 / YV1433 | *ulp1* | *ulp1::kanMX*<br>*YCpLac111-HIS3-ulp1-333 (HIS3)* | This study (c) |
| YV1451 | *CBC2-pA ulp1* | *CBC2-ProtA::His3 ulp1::kanMX*<br>*YCpLac111-ulp1-333 (LEU2)* | (Bretes et al., 2014) |
| YV1479 | *MLP2-pA* | *MLP2-ProtA::HIS3* | (Bretes et al., 2014) |
| YV1626 | *HEK2-3HA ulp1* | *HEK2-3HA::kanMX ulp1::kanMX*<br>*YCpLac111-HIS3-ulp1-333 (HIS3)* | This study * |
| YV1668 | *hek2Δ ulp1* | *hek2::kanMX ulp1::kanMX*<br>*YCpLac111-HIS3-ulp1-333 (HIS3)* | This study * |
| JR274 | *HEK2(K15R K29-30R)-3HA ulp1* | *hek2::kanMX ulp1::kanMX*<br>*YCpLac111-HIS3-ulp1-333 (HIS3)*<br>*pRS316-HEK2(K15R K29-30R)-3HA (URA3)* | This study * |
| YV1606 | *HEK2-3HA CBC2-pA* | *HEK2-3HA::kanMX CBC2-protA::HIS3* | This study * |
| YV1601 | *HEK2-3HA CBC2-pA ulp1* | *HEK2-3HA::kanMX CBC2-protA::HIS3*<br>*ulp1::kanMX*<br>*YCpLac111-ulp1-333 (LEU2)* | This study * |
| YV1756 | *HEK2-3HA MLP2-pA* | *HEK2-3HA::kanMX MLP2-protA::HIS3* | This study * |
| YV1757 | *HEK2-3HA MLP2-pA ulp1* | *HEK2-3HA::kanMX MLP2-protA::HIS3*<br>*ulp1::kanMX YCpLac111-ulp1-333 (HIS3)* | This study * |
| JR153 | *HEK2-pA ulp1* | *hek2::kanMX ulp1::kanMX*<br>*YCpLac111-HIS3-ulp1-333 (HIS3)*<br>*pRS316-HEK2-protA* | This study * |
| SWY518 | *wt* | **(W303)** | (Strawn et al., 2004) |
| SWY2950 | *nup145ΔFG nup57ΔFG*<br>*nup100ΔFG* | **(W303)** *myc-LoxP-nup145ΔGLFG*<br>*myc-LoxP-nup57ΔGLFG*<br>*HA-LoxP-nup100ΔGLFG* | (Strawn et al., 2004) |
| SWY2980 | *nup145ΔFG nup100ΔFG*<br>*nsp1ΔFGΔFxFG* | **(W303)** *myc-LoxP-nup145ΔGLFG*<br>*HA-LoxP-nup100ΔGLFG*<br>*Flag-LoxP-nsp1ΔFGΔFxFG* | (Strawn et al., 2004) |
| SWY2922 | *nsp1ΔFxFG-ΔFG* | **(W303)** *Flag-LoxP-nsp1ΔFxFG-ΔFG* | (Strawn et al., 2004) |
| SWY2801 | *nup1ΔFxFG* | **(W303)** *T7-LoxP-nup1ΔFxFG* | (Strawn et al., 2004) |
| YV929 | *ULP1-GFP mat a* | *Ulp1-GFP::HIS3MX* | Invitrogen |
| YV2049 | *ULP1-GFP* | **(W303)** *Ulp1-GFP::HIS3MX* | This study (d) |
| YV2052 | *ULP1-GFP nup145ΔFG*<br>*nup57ΔFG nup100ΔFG* | **(W303)** *Ulp1-GFP::HIS3MX*<br>*myc-LoxP-nup145ΔGLFG*<br>*myc-LoxP-nup57ΔGLFG*<br>*HA-LoxP-nup100ΔGLFG* | This study (d) |
| YV2050 | *ULP1-GFP nup145ΔFG*<br>*nup100ΔFG nsp1ΔFGΔFxFG* | **(W303)** *Ulp1-GFP::HIS3MX*<br>*myc-LoxP-nup145ΔGLFG*<br>*HA-LoxP-nup100ΔGLFG*<br>*Flag-LoxP-nsp1ΔFGΔFxFG* | This study (d) |
| YV2069 | *ULP1-GFP nsp1ΔFGΔFxFG* | **(W303)** *Ulp1-GFP::HIS3MX*<br>*Flag-LoxP-nsp1ΔFxFG-ΔFG* | This study (d) |
| YV2066 | *ULP1-GFP nup1ΔFxFG* | **(W303)** *Ulp1-GFP::HIS3MX*<br>*T7-LoxP-nup1ΔFxFG* | This study (d) |

Homozygous and heterozygous deletion strains were obtained from the Euroscarf deletion collection (www.euroscarf.de). GFP-tagged strains were purchased from Invitrogen.
\* obtained by transformation and/or successive crosses.
a. *HEK2* was deleted by homologous recombination with a cassette amplified from pFA6a-kanMX6.
b. *HEK2* was C-terminally tagged with 3 HA repeats by homologous recombination with a cassette amplified from pFA6a-3HA-kanMX6.
c. Segregant of a heterozygous diploid *ulp1::kanMX/ULP1+* transformed with the *YCpLac111-HIS3-ulp1-333*.
d. *ULP1* was C-terminally tagged with GFP by homologous recombination with a cassette amplified from the *ULP1-GFP* strain from the GFP collection.

**Table S2:**
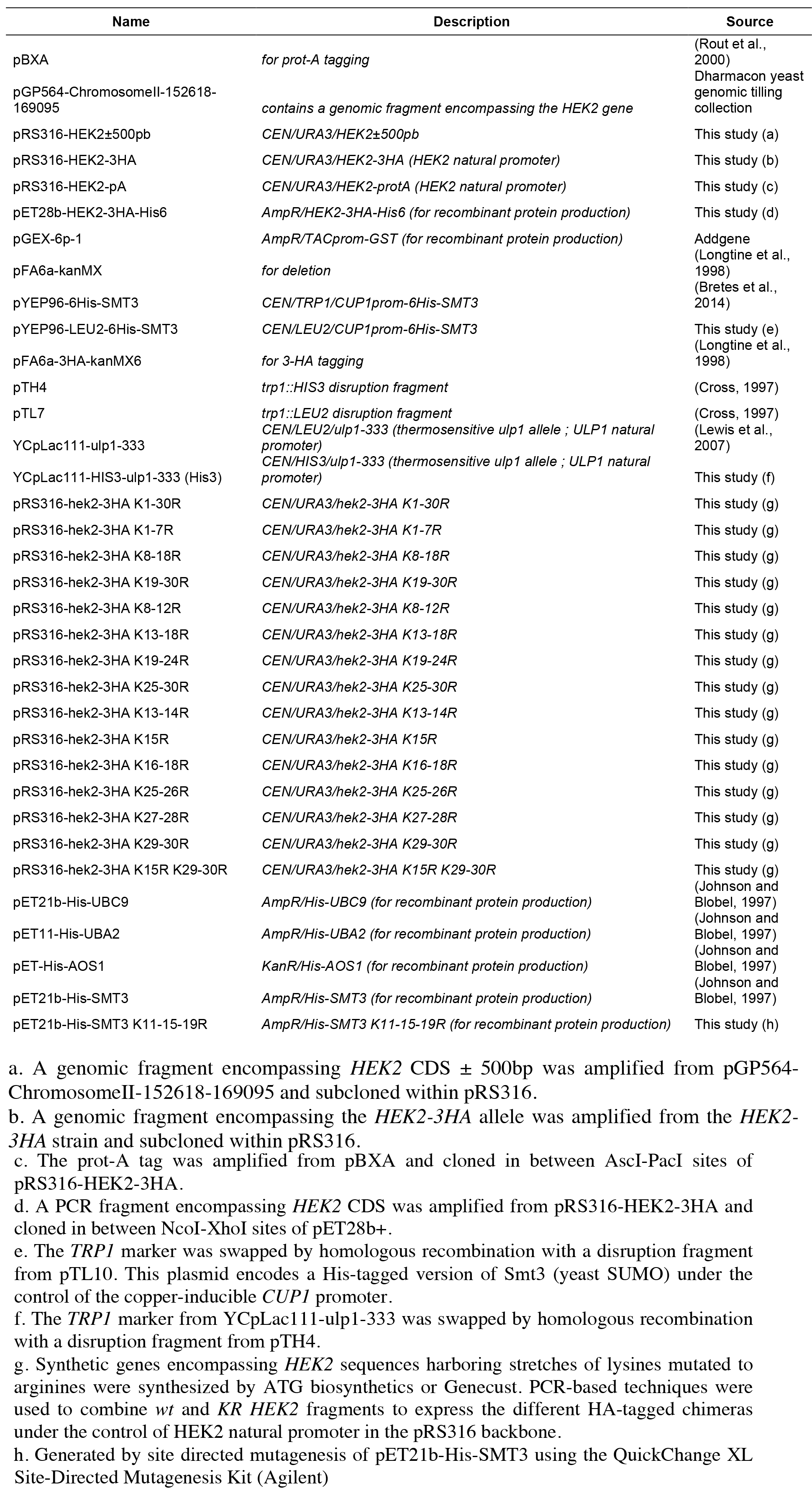
Plasmids used in this study

**Table S3:**
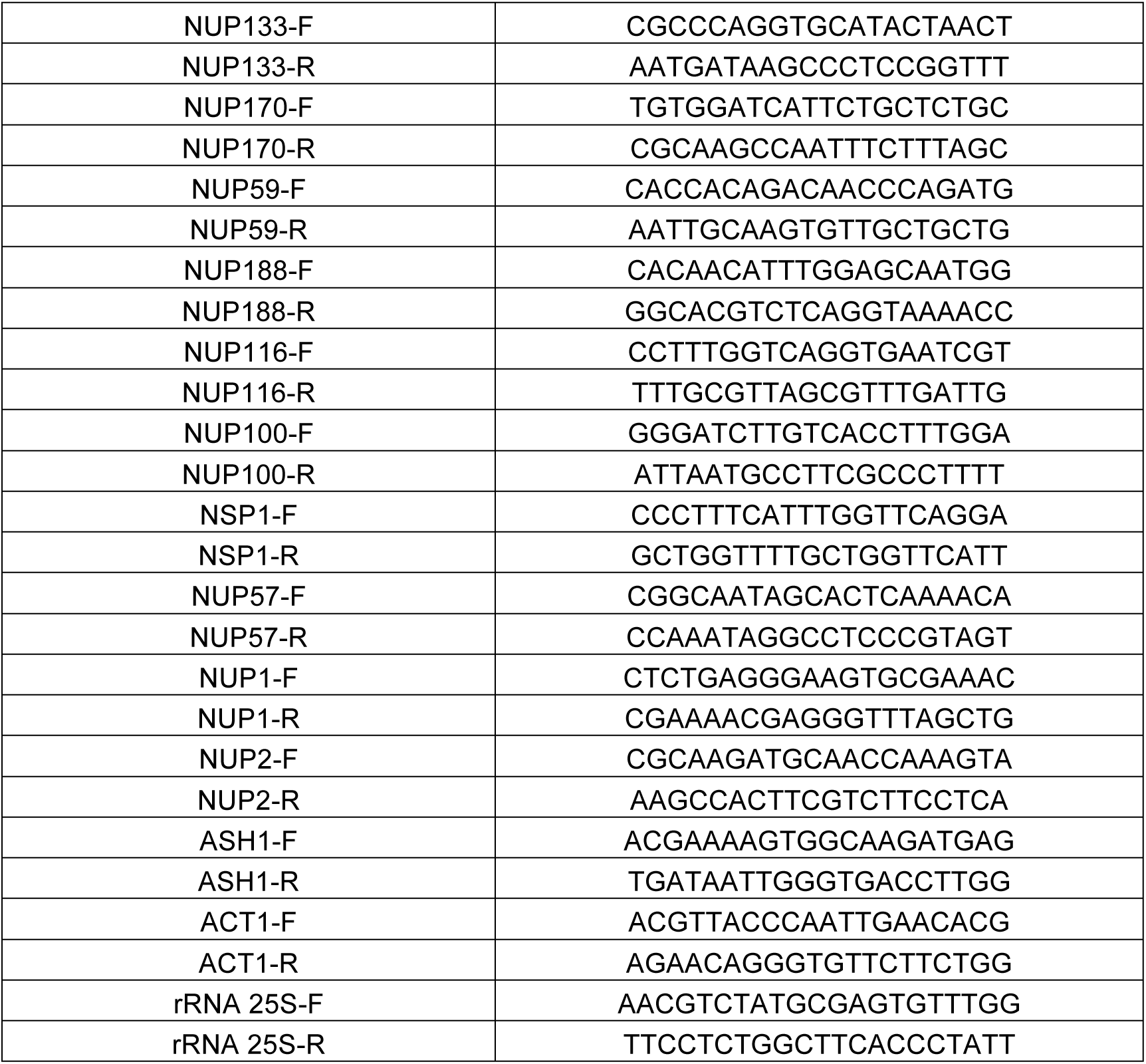
qPCR primers used in this study

**Figure S1.**
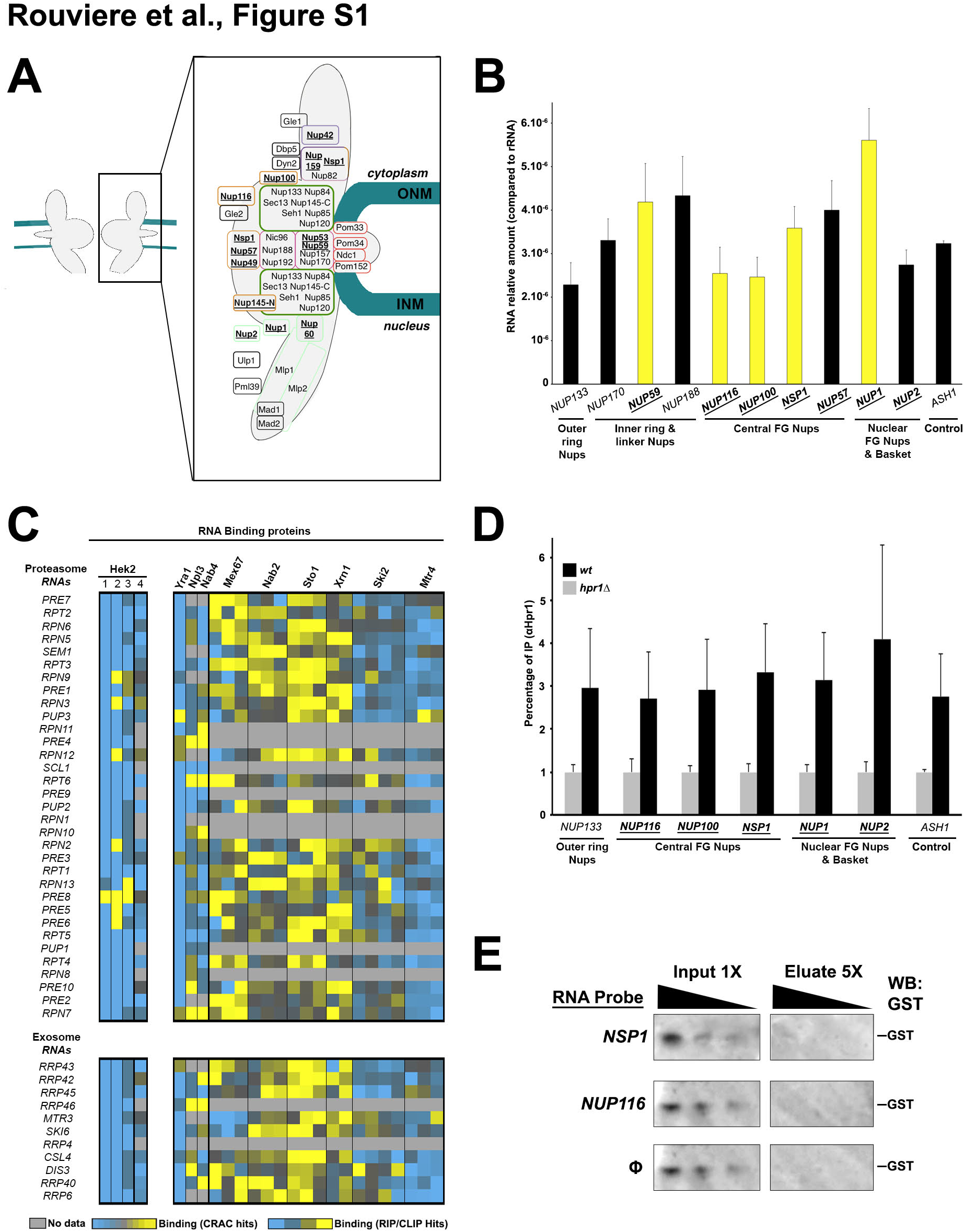
– related to Fig. 1. The association of Hek2 to a subset of *NPC* mRNAs is specific. **A,** Schematic representation of the yeast nuclear pore complex indicating the relative position of each nucleoporin within subcomplexes or along the NPC axis. FG-Nups appear in bold, underlined. ONM, outer nuclear membrane. INM, inner nuclear membrane. **B,** *NPC* mRNAs levels (mean ± SD; n=3; relative to rRNA) were measured by RT-qPCR in *HEK2-pA* strains. Hek2-bound *NPC* mRNAs appear in yellow. **C,** RBP binding was analyzed as in **Fig. 1A** for mRNAs encoding proteasome or exosome subunits. **D**, Hpr1-associated mRNAs were immunopurified using anti-Hpr1 antibodies (Bretes et al., 2014) and quantified by RT-qPCR using specific primer pairs. Percentages of IP (mean ± SD; n=3) are the ratios between purified and input RNAs, set to 1 for *hpr1Δ* control cells. **E**, Recombinant GST was incubated with streptavidin beads either naïve (Φ) or previously coated with biotinylated RNA probes encompassing Hek2-binding sites from *NSP1* or *NUP116*. Decreasing amounts of input and eluate fractions were loaded to allow comparison as in **Fig. 1E**.

**Figure S2.**
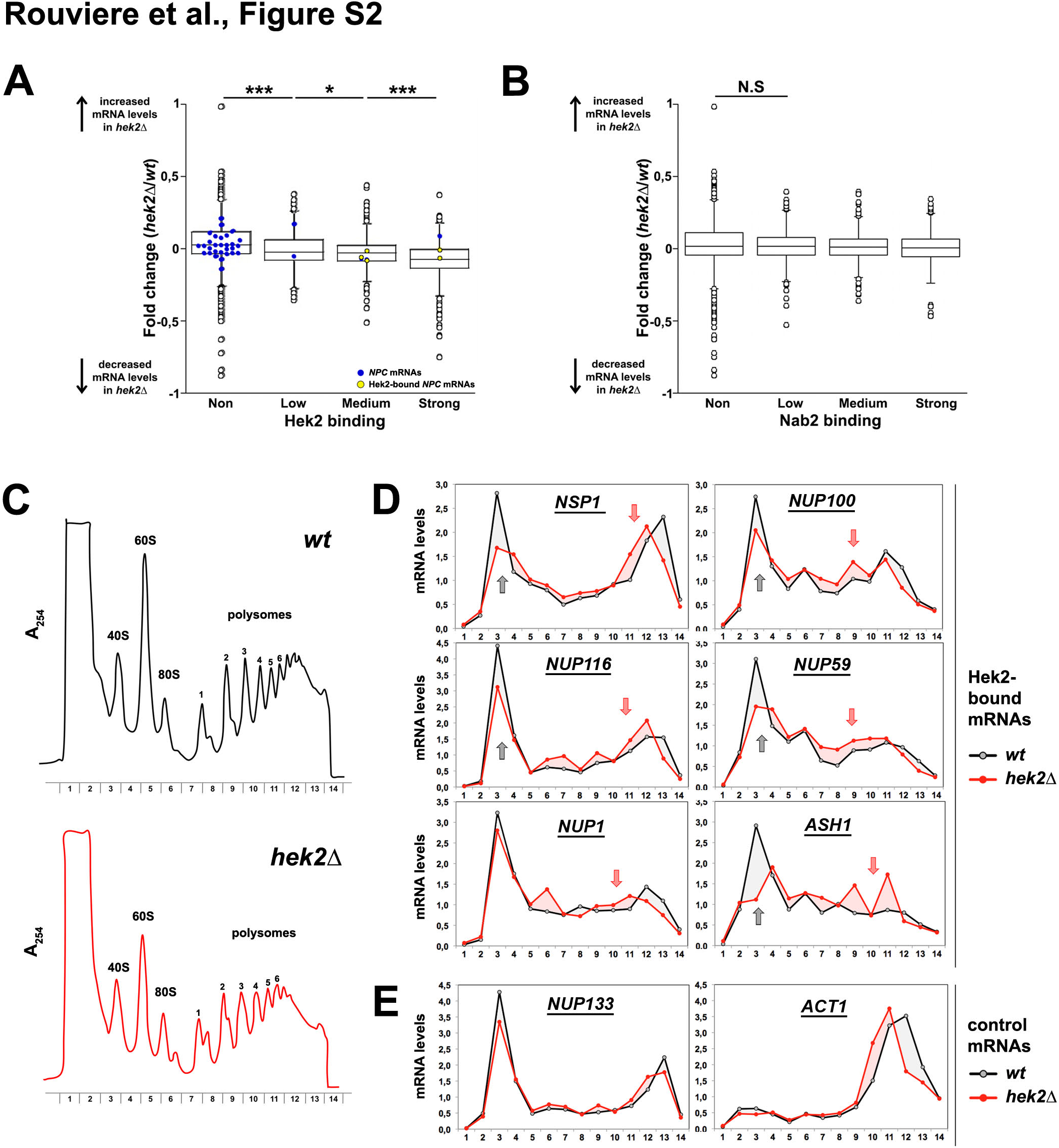
– related to Fig. 2. Hek2 binding does not affects the levels of *NPC* mRNA but rather modulates their ribosome occupancy. **A,** mRNAs were split in four categories depending on their binding to Hek2 (Hasegawa et al., 2008). For each group of transcripts, the averaged log2 of the *hek2Δ/wt* ratios calculated from two independent microarray hybridizations were plotted. mRNAs encoding NPCs components are highlighted in two different colors depending on their association to Hek2. Note that mRNAs strongly bound by Hek2 tend to be less abundant in the absence of this protein. * P&#x003C;0,5; ***P&#x003C;0,001 (Mann Whitney Wilcoxon test). **B,** The same analysis as in **A.** was performed after grouping the transcripts according to their binding to Nab2 (Kim Guisbert et al., 2005). Note the absence of correlation between Nab2 binding and the changes in mRNA levels scored upon *HEK2* inactivation. N.S: not significant. **C,** Polysome fractionation from *wt* and *hek2Δ* cells from the BY4742 background. The absorbance at 254nm (A_254_) recorded during the collection of the different fractions of the sucrose gradient is displayed. The positions of 40S, 60S, 80S ribosomal species and polysomes are indicated, as well as the number of ribosomes per mRNA in the polysomes fractions. Note that polysome profiles from these *hek2Δ* mutant cells exhibit reproducible discontinuities typical of half-mer formation, i.e. polysomes lacking stoichiometric amounts of both 60S and 40S ribosomal subunits. While this phenotype could reflect impaired 60S biogenesis, defective coupling of 60S subunits to 40S-mRNA complexes or general translational derepression (Cridge et al., 2010; Eisinger et al., 1997; Li et al., 2009), it was not observed in *hek2Δ* mutant cells of an alternate genetic background (W303, **Fig. 2C**), suggesting that it is not solely caused by *HEK2* inactivation. **D**, Relative distribution of the *NSP1*, *NUP100*, *NUP116*, *NUP59*, *NUP1* and *ASH1* mRNAs in polysome gradients from the same *wt* (black lines) and *hekΔ* (red lines) cells. mRNAs amounts in each fraction were quantified by RT-qPCR, normalized to the sum of the fractions and to the distribution of a control spike RNA. Grey arrows indicate a decrease in the amounts of mRNAs found in the light fractions in *hek2Δ* cells. Red arrows point to an increase in the quantity of mRNAs found in the polysomes fractions of the mutant. These results are representative of four independent experiments (two performed in the W303 background, two in the BY4742 background; see **Fig. 2**). **E,** Same as **D,** for *NUP133* and *ACT1* control mRNAs.

**Figure S3.**
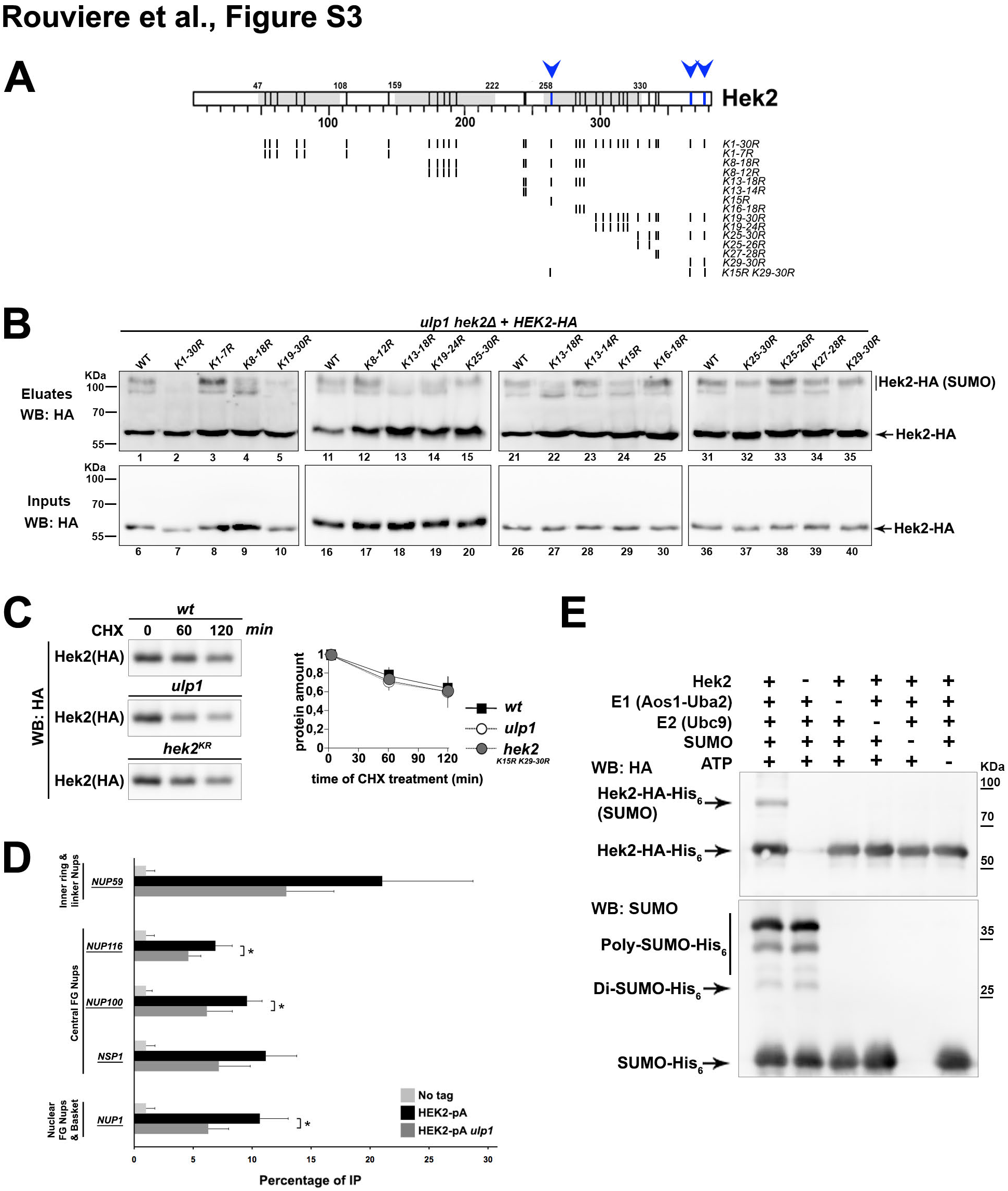
– related to Fig. 3. Characterization of Hek2 sumoylation. **A**, Schematic representation of the Hek2 protein and of the different *KR* mutants used in this study. Each vertical bar corresponds to a lysine residue and the KH-domains are displayed in grey, together with their boundaries as small numbers. For *KR* mutants, vertical bars represent the lysines that were mutated into arginines. The sumoylated residues identified in this study are indicated by blue bars and arrowheads. **B**, Hek2 sumoylation was analyzed in the indicated *KR* mutants as in **Fig. 3A-D**. Total lysates (”inputs”, *bottom panel*) and purified SUMO-conjugates (”eluates”, *top panel*) were analyzed by immunoblotting with anti-HA antibodies. The positions of the sumoylated and unmodified versions of Hek2-HA, as well as molecular weights, are indicated. **C**, Protein levels of HA-tagged versions of Hek2 were evaluated in *wt*, *ulp1* and *hek2 K15R K29-30R* (*hek2^KR^*) cells treated with cycloheximide (CHX) for the indicated time (minutes). Whole cell extracts were analyzed by western blotting using anti-HA antibody. The relative amounts of Hek2-HA (mean ± SD; n=2) were quantified over the time following CHX treatment and are expressed relative to t=0. **D**, Hek2-pA-associated mRNAs were immunopurified and quantified from *wt* (”no tag”), *HEK2-pA* and *HEK2-pA ulp1* cells as in **Fig. 1B**. Percentages of IP (mean ± SD; n=3) are the ratios between purified and input RNAs, further normalized to the amount of purified bait and set to 1 for the “no tag”. **E**, *In vitro* sumoylation of recombinant Hek2 was performed in the presence or the absence of the indicated components (”+” or “-”) and the reactions were analyzed by western blotting using anti-HA and anti-SUMO antibodies. The position of the sumoylated and unmodified versions of Hek2, of different poly-SUMO chains and of molecular weights are indicated. Note that the modified version of Hek2 is only detectable upon incubation of the recombinant protein with the unique combination of purified E1, E2, SUMO and ATP.

**Figure S4.**
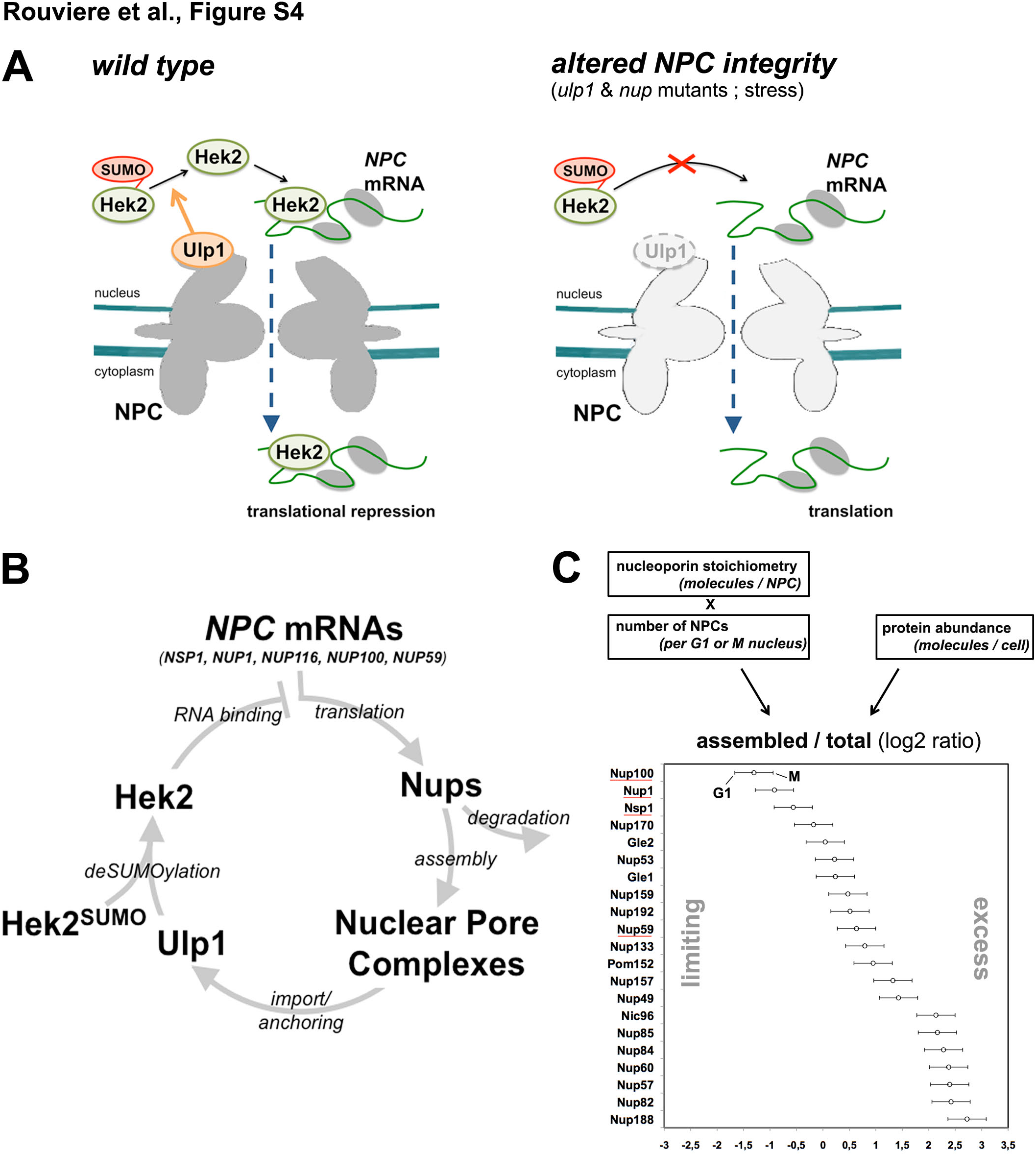
– related to Discussion. A SUMO-dependent feedback loop regulates the availability of nucleoporins. **A,** Schematic representation of the relationships between Ulp1 activity, Hek2 function and *NPC* mRNA expression scored in this study. In conditions of altered NPC integrity (right panel), decreased Ulp1 stability leads to the accumulation of sumoylated, inactive versions of Hek2, potentially releasing *NPC* mRNAs from their translationally-repressed state. **B,** Model for a feedback loop involving Ulp1 as a sensor of NPC integrity and controlling nucleoporin homeostasis through Hek2-mediated translational repression. **C,** For each nucleoporin, the amounts of proteins expected to be assembled in NPCs were calculated by multiplying the empiric values for NPC stoichiometry (Alber et al., 2007) by the total numbers of NPCs per nucleus, as counted in G1 or M cells (Winey et al., 1997). These values were further divided by the total cellular amounts of nucleoporins (as experimentally determined (Ghaemmaghami et al., 2003) and the log2 of these ratios were displayed. Horizontal bars reflect the expected variation between the G1 and M phases of the cell cycle. The more the displayed values are elevated, the more the corresponding nucleoporins are expected to be in excess as compared to the actual number of NPCs. Nucleoporins whose mRNAs are regulated by Hek2 are underlined in red. Note that this analysis does not include the nucleoporins which are part of other cellular complexes (i.e. Ndc1, Sec13, Seh1) or those for which abundances data were not available (Nup116, Nup145, Nup120, Nup2, Nup42, Pom34).

